# *Klebsiella pneumoniae* L-Fucose metabolism promotes gastrointestinal colonization and modulates its virulence determinants

**DOI:** 10.1101/2022.05.18.492588

**Authors:** Andrew W. Hudson, Andrew J. Barnes, Andrew S. Bray, M. Ammar Zafar

**Affiliations:** Department of Microbiology and Immunology, Wake Forest School of Medicine, Winston-Salem, NC 27101.

## Abstract

Colonization of the gastrointestinal (GI) tract by *Klebsiella pneumoniae* (*K. pneumoniae*) is generally considered asymptomatic. However, gut colonization allows *K. pneumoniae* to either translocate to sterile site within the same host or transmit through the fecal-oral route to another host. *K. pneumoniae* gut colonization is poorly understood, but knowledge of this first step toward infection and spread is critical for combatting its disease manifestations. *K. pneumoniae* must overcome colonization resistance (CR) provided by the host microbiota to establish itself within the gut. One such mechanism of CR is through nutrient competition. Pathogens that metabolizes a broad range of substrates have the ability to bypass nutrient competition and overcome CR. Herein, we demonstrate that in response to mucin derived fucose, the conserved fucose metabolism operon (*fuc*) of *K. pneumoniae* is upregulated in the murine gut and subsequently show that fucose metabolism promotes robust gut colonization. Growth studies using cecal filtrate as a proxy for the gut lumen illustrates the growth advantage that the *fuc* operon provides *K. pneumoniae*. We further show that fucose metabolism allows *K. pneumoniae* to be competitive with a commensal *E. coli* isolate (Nissle). However, Nissle is eventually able to out-compete *K. pneumoniae*, suggesting that it can be utilized to enhance CR. Lastly, we observed that fucose metabolism positively modulates hypermucoviscosity, auto-aggregation, and biofilm formation, but not capsule biogenesis. Together, these insights enhance our understanding of the role of alternative carbon sources on *K. pneumoniae* gut colonization and the complex relationship between metabolism and virulence in this species.

**Importance:** *Klebsiella pneumoniae* (*K. pneumoniae*) is a leading cause of hospital-acquired infection. Treatment of infection by *K. pneumoniae* isolates is becoming difficult as this pathogen becomes increasingly antibiotic resistant. While there has been extensive investigation into the disease states associated with *K. pneumoniae*, its colonization of the gastro-intestinal (GI) tract is poorly understood. Epidemiological data suggest that in many cases the strain that colonizes the GI tract causes disease manifestations in the same host. Herein, we used our newly developed murine model of *K. pneumoniae* gut colonization, where colonization is achieved without disrupting the resident gut microbiota. We demonstrate that *K. pneumoniae* uses fucose as an alternative carbon source present in the gut lumen to overcome the intense nutritional competition. We further illustrate that *K. pneumoniae,* through fucose metabolism, is initially competitive with the probiotic *E coli* isolate Nissle 1917 (EcN). Lastly, we show that fucose metabolism modulates several virulence determinants of *K. pneumoniae*. Thus, our results provide new insight into the role fucose metabolism plays in gut colonization and virulence of *K. pneumoniae*, and furthermore identify EcN as having the ability to out-compete *K. pneumoniae* and be used as a probiotic.

## Introduction

The human gastrointestinal (GI) tract is a competitive environment in which native microbiota contend with each other and host cells for nutrients (1, 2). This intense pressure for nutrient acquisition is one mechanism through which the host and the gut microbiota deter an incoming pathogen from carving out its metabolic niche in the GI tract - a term known as colonization resistance (CR). In turn, the ability to bypass nutrient competition allows an incoming microorganism to overcome CR (2-4).

*Klebsiella pneumoniae* (*K. pneumoniae*), a medically relevant Gram-negative bacterium, has gained notoriety with the emergence of multi-drug resistant strains and is considered a priority pathogen by WHO for which urgent new therapeutics are required (5). Studies have shown that the *K. pneumoniae* isolate that colonizes the GI tract can causes disease manifestation within the same host (6-8). Disease states associated with *K. pneumoniae* include pneumonia, bacteremia, pyogenic liver abscesses, and urinary tract infections (UTIs) (9, 10). Epidemiological data suggest that *K. pneumoniae* spreads from the gut by translocating to sterile sites within the host and transmitting through the fecal-oral route from one host to another (6, 8, 11). Despite CR, *K. pneumoniae* forms part of the healthy human GI microbiota (9, 12). However, unlike enterohemorrhagic *Escherichia coli* (EHEC) and *Salmonella enterica* ssp. Typhimurium (*S. typhimurium*), which modify the gut and induce a robust inflammatory response (13, 14), *K. pneumoniae* colonizes asymptomatically and uses the gut as a silent reservoir for infection and transmission (6, 9-11).

Despite the clinical importance of *K. pneumoniae*, how it colonizes the gut is poorly understood (10). Although a few recent studies have focused on gut commensals that provide CR against *K. pneumoniae* (15, 16). Metabolic flexibility is critical for bypassing nutritional limitations in the GI tract to overcome CR. Commensal *Escherichia coli* (*E. coli*) can metabolize a broad range of carbon sources in the gut (17). For *E. coli*, carbon sources exist on a hierarchy of “preference” depending on energetic efficiency, with more preferred carbon sources such as glucose having a more significant impact on colonization than others (18). Similarly, *K. pneumoniae* possesses a diverse set of genes for carbohydrate metabolism (19), and consequently, it may co-metabolize multiple carbon sources to colonize the gut and initiate its unique infectious cycle.

The available sugars in the GI tract include those that are derived from mucins, heavily glycosylated proteins which serve as the primary constituents of the gut mucus layer (20, 21). *Bacteroides* species produce hydrolytic enzymes including fucosidases that cleave mucin-associated fucose to gain access to energetically rich core glycans (20, 22, 23). Many commensals and pathogens alike cannot cleave mucin but can metabolize the liberated sugar moieties including fucose (18, 22, 23). Despite being a carbon source of low preference for *E. coli* (18), fucose utilization gives EHEC a competitive advantage in the gut (17) and *S. Typhimurium* was shown to rely on fucose and sialic acid liberated by *Bacteroides thetaiotaomicron* (*B.t*) to colonize the murine GI tract (22).

Moreover, *S. Typhimurium* induces fucosylation in the small intestine and uses fucosylated mucin as a binding site for Std fimbriae, enhancing its ability to colonize and cause disease (24, 25). The *K. pneumoniae* genome encodes a highly conserved fucose metabolism operon (*fuc*), whose role in its pathogenesis remains largely unexplored. An *in vivo* transposon-directed insertion sequencing (TraDIS) screen using *Galleria mellonella* identified several genes of the *fuc* regulon to be critical for *K. pneumoniae* infection (26). Together, these data suggest a role for fucose metabolism in the ability of *K. pneumoniae* to overcome CR and colonize the GI tract.

Metabolites not only facilitate pathogen establishment in the host but can also modulate expression of virulence genes (27-29). For instance, EHEC uses exogenous fucose as a signal to regulate its virulence expression when colonizing the gut (27). In addition, nutrient metabolism is a known modulator of virulence determinants in *K. pneumoniae* (10, 29, 30), though as with all aspects of *K. pneumoniae* GI colonization, virulence modulation in the gut is still poorly understood.

Historically, mouse models of *K. pneumoniae* gut colonization have relied on antibiotic pre-treatment that depletes the resident gut microbiota and reduces CR (12, 31). Consequently, a decline in microbiota complexity allows *K. pneumoniae* to overcome CR, but it hinders our ability to provide a molecular understanding of pathogen-microbiome interactions. Our newly developed murine model for *K. pneumoniae* GI colonization alleviates these concerns, preserving the intact microbiome and associated CR to recapitulate natural colonization of the healthy human gut (12). Herein, use of our murine model revealed the importance of fucose utilization in overcoming CR by a pathogen that silently colonizes the GI tract. Analysis of complimentary *in vitro* assays provided a molecular understanding of the contribution of different carbon sources on known virulence gene expression of *K. pneumoniae*. In this work, we advance the understanding of the metabolic dynamics that underlie this critical stage of *K. pneumoniae* pathogenesis.

## Results

### *K. pneumoniae* uses fucose as an alternative carbon source

The ability to metabolize alternative carbon sources is one molecular mechanism that allows enteric pathogens to overcome CR (32). The *fuc* operon, conserved in many Enterobacteriaceae, allows the metabolism of fucose that has been liberated from glycosylated mucin proteins by resident gut microbiota (20, 22). As *K. pneumoniae* possesses a conserved *fuc* operon that has not been previously characterized (**Fig. 1A**), we hypothesized that it metabolizes fucose as an alternative carbon source to colonize the gut.

**Fig 1.**
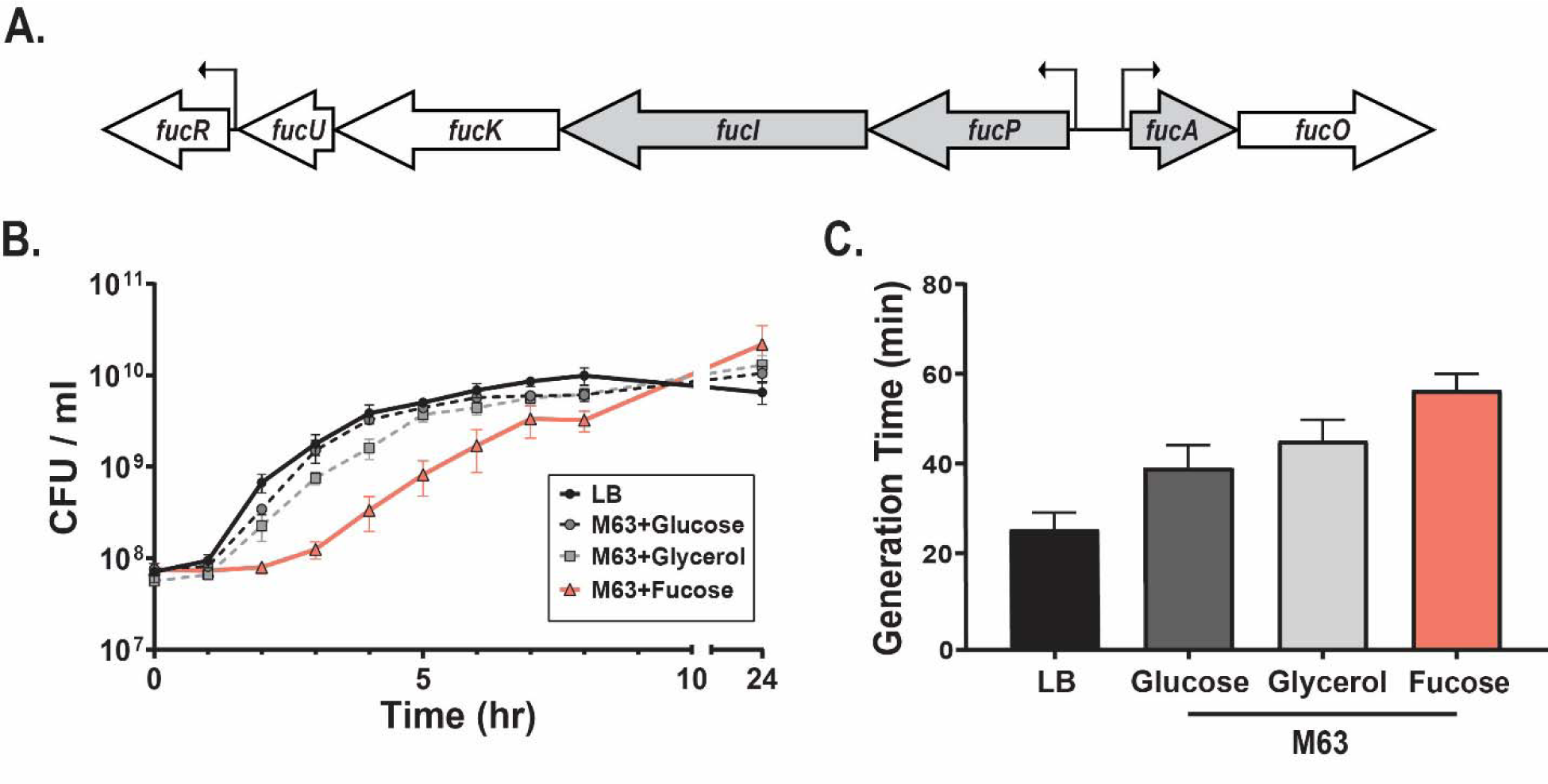
*K. pneumoniae* uses fucose as an alternative carbon source. **(A)** Schematic of the *fuc* operon of the *K. pneumoniae* isolate KPPR1S, with genes used for qRT-PCR highlighted. **(B)** Growth kinetics as depicted by CFU/ml over time and **(C)** mid-log phase generation time of *K. pneumoniae* in LB-Lennox and in M63 minimal media (MM) with either 0.5% glucose, glycerol, or fucose as the sole carbon source. Mean ± SEM for ≥3 independent experiments is shown.

To determine whether *K. pneumoniae* can metabolize fucose, we grew our model clinical strain KPPR1S (derivative of ATCC 43816; WT) (33, 34) in M63 minimal media (MM) supplemented with fucose as the sole carbon source. As observed, *K. pneumoniae* grew in the presence of fucose (**Fig. 1B**). Carbon sources are preferentially utilized hierarchically depending on energetic efficiency, with glucose being the most preferred carbon source in many species (17, 18, 35). In comparison to rich media (lysogeny broth [LB]) and MM supplemented with either glucose or glycerol, *K. pneumoniae* had slower growth kinetics when grown on fucose (**Fig. 1B**). Moreover, generation time, the time for a bacterial population to double during logarithmic growth, was higher when fucose was provided as the sole carbon source, indicating slower growth (**Fig. 1C**). Thus, while *K. pneumoniae* can metabolize fucose as its sole carbon source, it is a less energetically efficient alternative to glucose and glycerol.

### *K. pneumoniae fuc* operon is upregulated in the presence of fucose

To determine if the *K. pneumoniae fuc* operon is transcribed, we performed quantitative Reverse-Transcription PCR (qRT-PCR) on RNA isolated from bacterial samples grown in MM supplemented with either glucose or fucose. We examined expression of the *fuc* operon by directing primers at three *fuc* operon genes: fucose permease (*fucP*), which imports fucose into the cell; fucose isomerase (*fucI*), an early-acting enzyme in fucose metabolism; and fucose aldolase (*fucA*), an enzyme that acts late in fucose metabolism. All three genes were strongly upregulated when *K. pneumoniae* was grown in the presence of fucose compared to glucose (**Fig. 2A**) and revealed the tight regulation of this operon by carbon availability.

**Fig 2.**
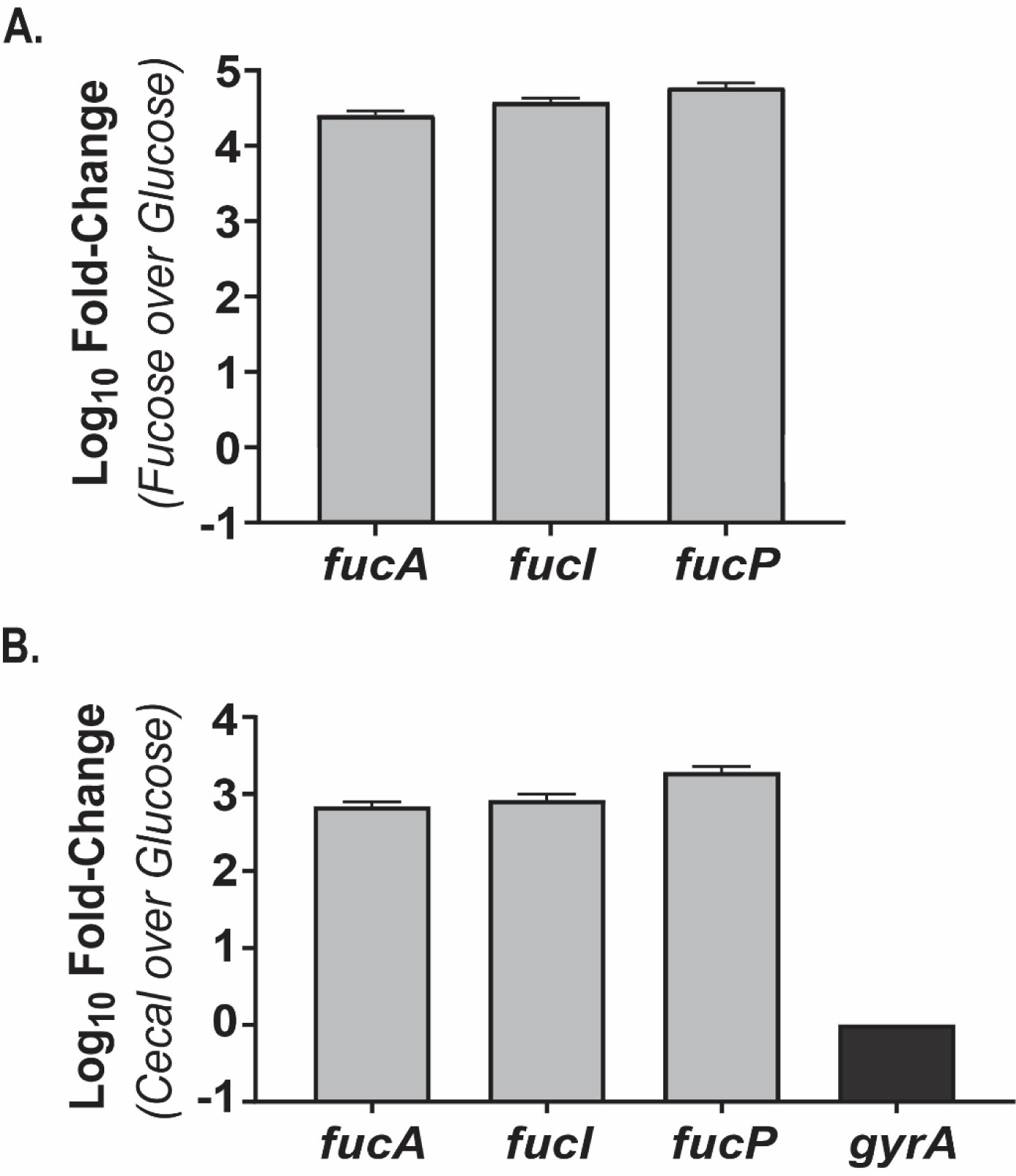
*K. pneumoniae fuc* operon is upregulated in the presence of fucose and in the murine GI tract. **(A)** Quantitative Reverse-Transcription PCR (qRT-PCR) comparing *fuc* gene expression of fucose- and glucose-grown *K. pneumoniae.* Shown is log_10_ of fold change in transcription of *fucA*, *fucI*, and *fucP* between *K. pneumoniae* grown in either MM + .5% fucose or grown in MM + .5% glucose. *gyrA* was used as the housekeeping gene for 2^-*ΔΔC*^*T* analysis. For each biological replicate (n ≥ 3 per condition), qRT-PCR was conducted in triplicate. **(B)** qRT-PCR showing *fuc* operon expression in the murine GI tract. Shown is fold-change in transcription of *fucA*, *fucI*, and *fucP* comparing RNA isolated from cecal contents from either *K. pneumoniae* infected mice or grown in MM + .5% glucose. Mice were infected by oral feeding of 10^6^ CFU, and cecal contents were harvested 15 days post-infection. KPPR1S *16s* (*rrsA*)-specific primers were used as the housekeeping gene for 2*^-ΔΔC^T* analysis. The values were further normalized to *gyrA* expression. For each biological replicate (n ≥ 3 for *in vitro* and *in vivo* samples), qRT-PCR was conducted in triplicate. Shown is mean ± SEM.

Next, to determine if *K. pneumoniae* encounters fucose in the murine gut, we employed our murine model of *K. pneumoniae* GI tract colonization (12). We observed a strong upregulation of the *fuc* operon (**Fig. 2B**) from cecal RNA isolated from *K. pneumoniae* inoculated mice, suggesting that it encounters and responds to fucose in the gut lumen. Collectively, our data suggest there is an increase in the transcript level of *fuc* genes when *K. pneumoniae* is exposed to fucose under both *in vitro* and *in vivo* conditions.

### Fucose metabolism promotes gastrointestinal colonization of *K. pneumoniae*

To assess the contribution of fucose utilization on the ability of *K. pneumoniae* to colonize the gut, we created an isogenic *ΔfucI* mutant that is unable to metabolize fucose (**Fig. S1A**). Mice with an intact microbiota were orally inoculated with the wild-type (WT), *ΔfucI*, or *fucI* chromosomally reconstituted at the native site (*fucI*+). Fecal shedding was used as a non-invasive metric for daily colonization. For 15 days post-infection, mice inoculated with the *ΔfucI* strain generally shed poorly (indicating reduced colonization fitness), while the wild-type (WT) and *fucI*+ strains shed robustly (**Fig. 3A**). The shedding phenotype correlated with colonization of the GI tract (**Fig. S1B-C**). Two shedding populations were observed with mice inoculated with the *ΔfucI* strain; one that shed the mutant strain poorly and the other that shed as WT (**Fig. S1D**). We attribute the colonization variability of the *ΔfucI* strain to subtle differences in gut microbiota of individual mice (12, 36), which then manifests as either higher or lower CR threshold for the *ΔfucI* strain. Subsequently, we investigated whether the WT would rescue the gut colonization defect of *ΔfucI* with co-inoculation experiments. As observed through daily shedding and colonization density in the gut, the WT strain out-competed the *ΔfucI* mutant (**Fig. 3B-C**). Collectively, the *K. pneumoniae ΔfucI* strain has a significant fitness defect in gut colonization, suggesting that the ability to metabolize fucose is critical for consistent, robust GI colonization.

**Fig 3.**
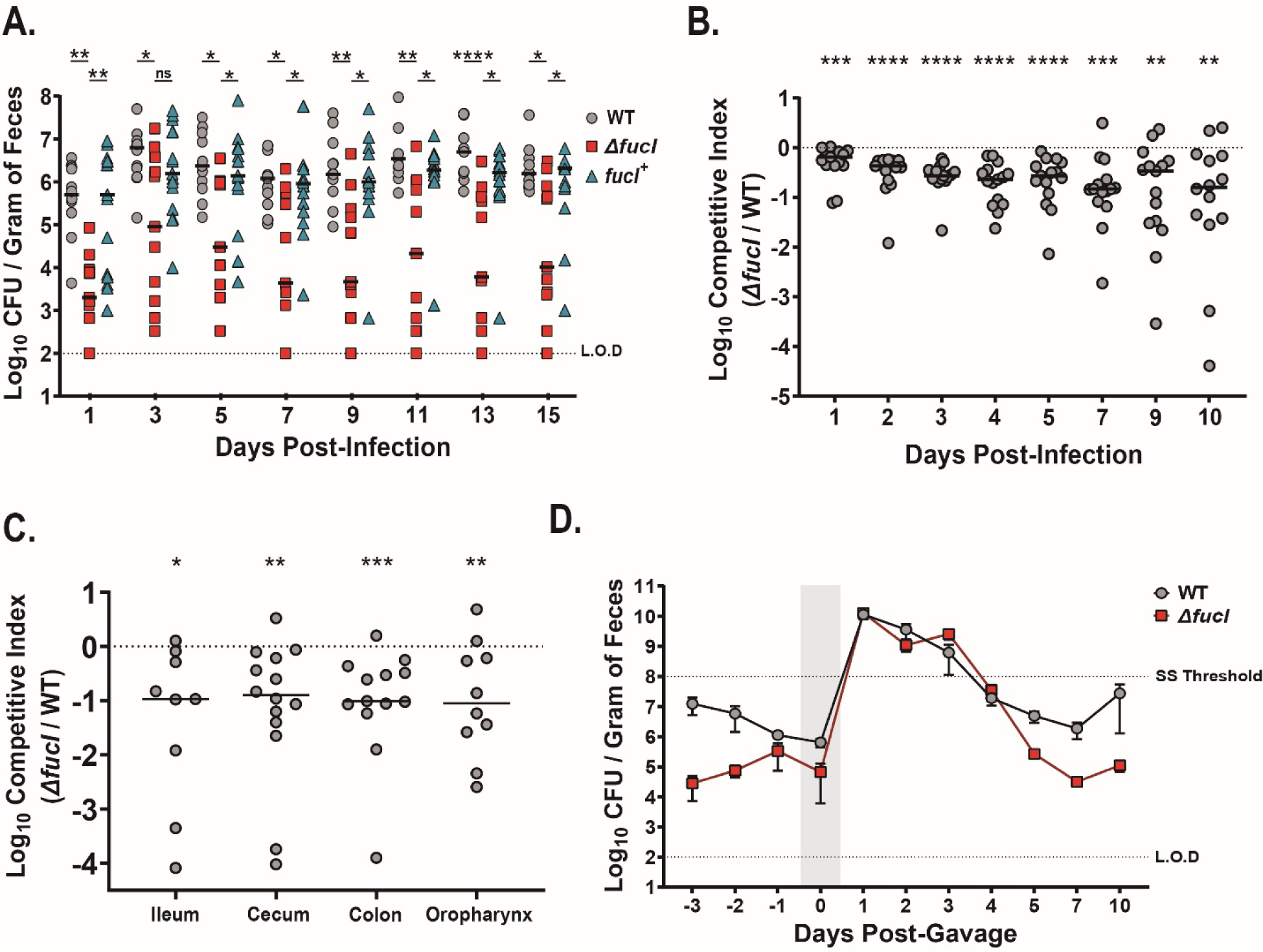
Fucose utilization by *K. pneumoniae* promotes gut colonization. **(A)** Fecal shedding of *K. pneumoniae* infected mice. Mice were orally inoculated with 10^6^ CFU of WT *K. pneumoniae*, the isogenic mutant (*ΔfucI)*, or the chromosomally complemented isolate (*fucI^+^)* (n ≥ 10 for each group). On indicated days, *K. pneumoniae* was enumerated from collected feces. Each symbol represents a single mouse on a given day, bars indicate median bacterial shedding, and the dashed line indicates limit of detection (L.O.D). A Kruskal-Wallis followed by Dunn’s test of multiple comparisons was performed at each time point. **(B)** Fecal shedding from *in vivo* competition experiment. Mice were orally infected with a 1:1 mixture of WT and *ΔfucI* strain and fecal shedding was enumerated on indicated days (n ≥ 10). The CI was determined as described in Materials and Methods. Each symbol represents a single mouse on a given day. Values above the dashed line indicate the mutant (Δ*fucI*) out-competing WT, whereas values below the dashed line indicate WT out-competing *ΔfucI*. **(C)** Colonization density at the end of the *in vivo* competition study. Colonization density of the WT and the Δ*fucI* strain in the ileum, cecum, colon, and oropharynx was enumerated and CI was calculated. **(B, C)** Bars indicate median value. Statistical significances of CIs were calculated using Wilcoxon signed-rank test with a theoretical median of 0. **(D)** Antibiotic treatment leads to development of supershedder state in mice inoculated with the WT or the *ΔfucI* strain. Shown is mean fecal shedding of mice orally inoculated with either the WT or the *ΔfucI* strain, with a single dose of streptomycin sulfate (5 mg/200 μl) administered by oral gavage 5 days post-infection (n = 5 for each strain, bars represent SEM). Top and bottom dashed lines indicate the supershedder threshold and the limit of detection, respectively. * P < 0.05, ** P < 0.01, *** P < 0.001, **** P < 0.0001

Our murine model of *K. pneumoniae* GI colonization preserves the native microbiome and therefore mimics colonization of a healthy gut through the fecal-oral route (12). However, many patients in a hospital setting are on antibiotics, which disrupt the native gut microbiota and thus reduce CR. Previous studies have shown that disruption of host-microbiota by antibiotics manifests as a transient “supershedder” state in which resistant enteric pathogens bloom to high levels in the gut, resulting in increased transmission (12, 37, 38). Additionally, fucose metabolism furnishes *S. Typhimurium* with a competitive advantage in the gut during antibiotic mediated gut dysbiosis (22). Hence, we assessed the fitness advantage to *K. pneumoniae* conferred by fucose metabolism immediately following antibiotic treatment. Mice mono-infected with either the WT or *ΔfucI* were gavaged with streptomycin to induce a temporary supershedder phenotype. The ability to metabolize fucose did not confer a colonization advantage during the supershedder state (**Fig. 3D**), suggesting that the observed effect of fucose utilization in the GI tract is dependent on nutritional competition. Furthermore, unlike *S. Typhimurium*, fucose metabolism is dispensable for *K. pneumoniae* following antibiotic treatment, as it potentially has access to more preferred carbon sources during post-antibiotic expansion.

### Fucose metabolism provides *K. pneumoniae* with a growth advantage

To investigate if fucose metabolism provides *K. pneumoniae* with a growth advantage in the GI tract, we performed growth studies in homogenized cecal filtrate (CF) to recapitulate conditions of the GI microenvironment *in vitro*. Filtration removes cells and tissue debris but retains small metabolites such as fucose. Under aerobic conditions, both the WT and the *ΔfucI* mutant replicated in CF (**Fig. S2A**), and the transcript abundance of the *fuc* genes increased in the WT, indicating metabolism of fucose occurred during growth in this medium (**Fig. S2B**). Although both strains grew, the *ΔfucI* mutant did not reach the same cell density as the WT at 24 hours post-inoculation, suggesting the inability to metabolize fucose present in the media caused a growth defect. Moreover, in competitive growth studies between the WT and the *ΔfucI* mutant, the WT consistently out-competed the mutant in CF (**Fig. 4A**). In contrast, this out-competition did not occur in glucose supplemented MM where fucose was absent (**Fig. 4A**). Our data suggest that fucose utilization confers a competitive advantage in CF, as in the mouse, demonstrating the fidelity of the CF growth media in recapitulating the metabolic conditions of the GI tract.

**Fig 4.**
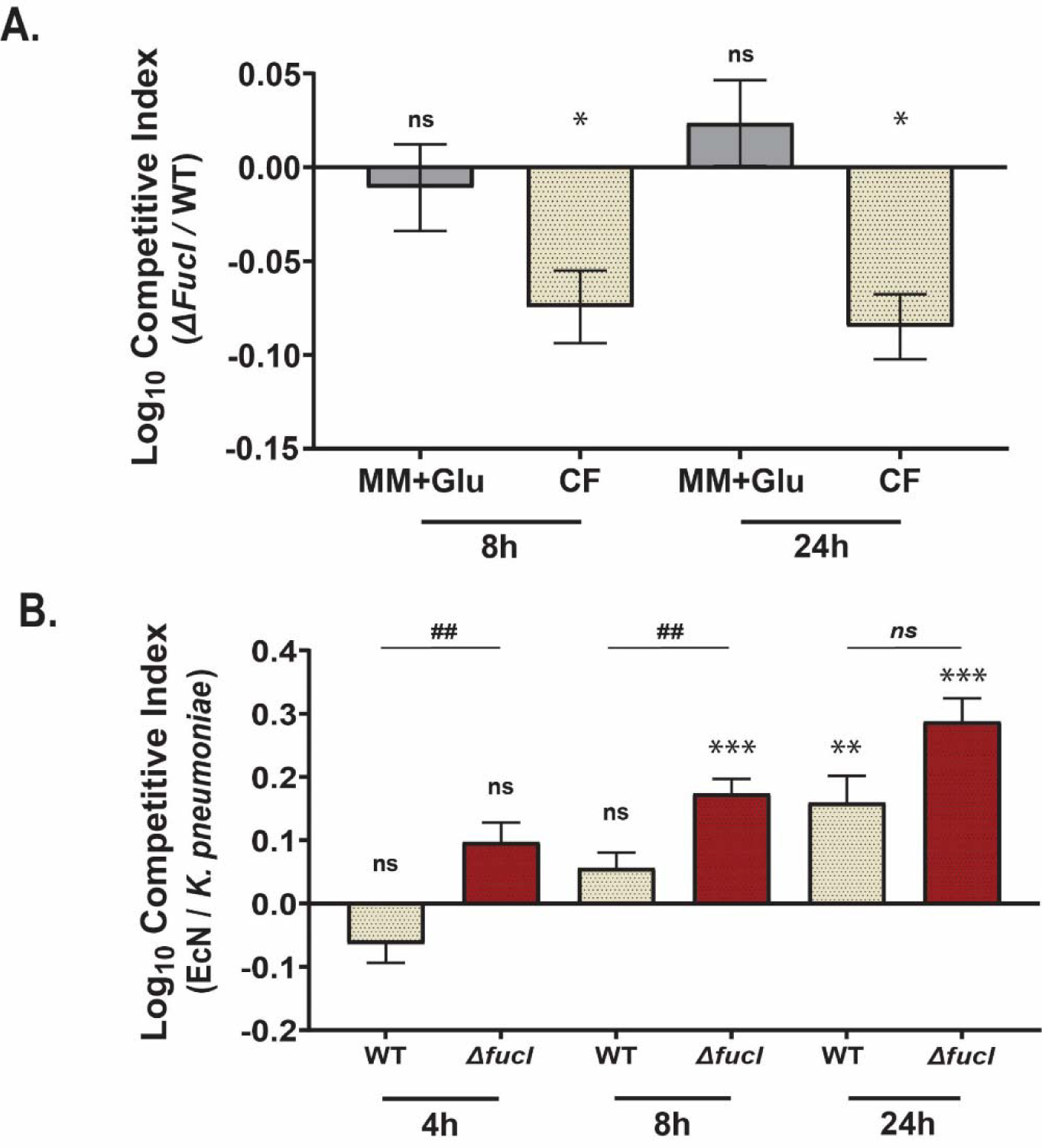
Fucose metabolism affords *K. pneumoniae* with a growth advantage. **(A)** *In vitro* competition experiments between the WT and the *ΔfucI K. pneumoniae*. Both strains were inoculated 1:1 into MM + .5% glucose or into cecal filtrate (CF). CFUs were enumerated and CI values determined at 8 and 24 hours post-inoculation. **(B)** CF competition experiments between *K. pneumoniae* and EcN. Either the WT or the *ΔfucI K. pneumoniae* were inoculated into CF with EcN at a 1:1 ratio. CFUs were enumerated at 4, 8 and 24 hours post-inoculation and CI values were calculated. Shown is mean ± SEM from ≥3 individual experiments. Statistical significances of CIs were calculated using Wilcoxon signed-rank test with a theoretical median of 0. * P < 0.05, ns - not significant. A Mann-Whitney *U* test was performed between the WT and the *ΔfucI* strain CIs at the indicated time points. ## P < 0.01, *ns* – not significant

We next used the probiotic *E. coli* strain Nissle 1917 (EcN) (39) as a representative human gut commensal to determine whether it could outcompete *K. pneumoniae*. We observed that EcN not only grew in CF (**Fig. S2A**), but out-competed both the WT and *ΔfucI* at 24 hours post-inoculation, mimicking natural commensal opposition in the gut (**Fig. 4B**). Furthermore, at 4 and 8 hours post-inoculation in CF, the *ΔfucI* strain but not the WT was out-competed by EcN (**Fig. 4B**). However, this effect was only seen in CF, as in glucose-supplemented MM, the WT and *ΔfucI* mutant out-competed EcN (**Fig. S3A**). Similar results were obtained with fucose-supplemented MM (**Fig. S3B**). Our *in vitro* growth studies in CF suggest that fucose metabolism provides the WT a competitive advantage against the *ΔfucI* mutant and that it enhances the ability of *K. pneumoniae* to compete with EcN.

### Fucose metabolism enhances hypermucoviscosity without affecting capsule amount

As metabolic conditions can modulate virulence phenotypes (27, 28), we explored whether fucose utilization impacts known virulence determinants in *K. pneumoniae*. Hypermucoviscosity (HMV) is a hallmark of hypervirulent *K. pneumoniae* strains and contributes to its virulence (9, 40). Historically, a colony string test has been used to characterize the HMV phenotype, though a recently developed low-speed centrifugation assay provides a more reliable quantification of HMV (41, 42). Furthermore, HMV phenotype is considered to be distinct from high capsule production (41, 43). Using the low-speed sedimentation assay, we quantified HMV when *K. pneumoniae* was grown in the presence of fucose. We observed high supernatant optical density (ODs), measured at 600 nm when *K. pneumoniae* was grown in rich media (**Fig. 5A**). In contrast, low supernatant ODs were observed when *K. pneumoniae* was grown in MM, independent of carbon source. Supplementing our MM with casamino acids allowed for better growth dynamics and a more robust HMV phenotype, allowing us to determine the contribution of carbon source on the HMV phenotype (**Fig. 5A**). Notably, compared to glucose and glycerol, MM supplemented with fucose led to a significant increase in mucoviscosity.

**Fig 5.**
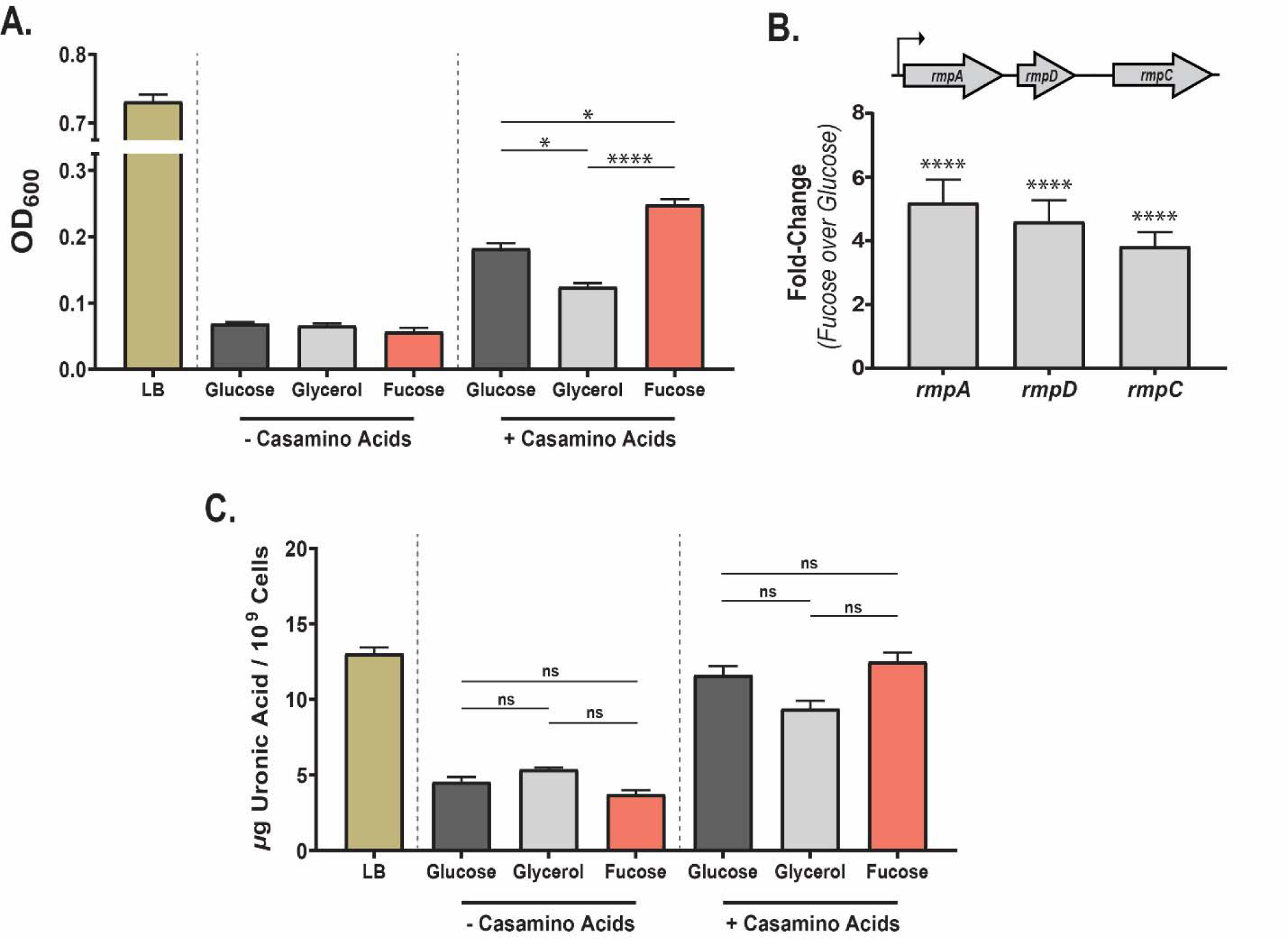
Fucose utilization positively modulates hypermucoviscosity, without impacting capsule levels. **(A)** Comparison of hypermucoviscosity (HMV) of *K. pneumoniae* grown in LB and in MM with either glucose, glycerol or fucose as the carbon source with or without the addition of casamino acids. **(B)** qRT-PCR comparing *rmp* locus gene expression from *K. pneumoniae* grown in MM with either glucose or fucose as the carbon source and addition of casamino acids. Shown is fold-change difference. *gyrA* was used as the housekeeping gene for 2*^-ΔΔC^T* analysis. For each biological replicate (n ≥ 3 per condition), qRT-PCR was conducted at least twice. **(C)** Uronic acid content assay to determine capsule amount was conducted on *K. pneumoniae* grown in LB and in MM described above. A Kruskal-Wallis followed by Dunn’s test of multiple comparisons was performed comparing HMV and uronic acid content of *K. pneumoniae* grown in M63 with different carbon sources with or without casamino acids. A Mann-Whitney *U* test was performed for each gene to determine significance. For all graphs, shown is mean ± SEM. * P < 0.05, **** P < 0.0001, ns - not significant

The *rmp* locus, specifically *rmpD*, was identified as required for the HMV phenotype (43). To determine whether the increase in HMV in fucose supplemented MM was a result of increased *rmpD* expression, we performed qRT-PCR on RNA isolated from *K. pneumoniae* grown in MM supplemented with casamino acids and either glucose or fucose. RNA transcripts of all genes in the *rmp* locus, including *rmpD*, were increased in cultures grown in fucose compared to glucose, suggesting the fucose-mediated increase in HMV was due to the upregulation of *rmpD* (∼4 fold) (**Fig. 5B**).

We next determined whether fucose utilization modulated capsular polysaccharide (CPS). CPS is considered one of the main virulence factors of *K. pneumoniae* and protects against host mediated clearance (10, 44), and nutrient conditions, including the carbon source, have been shown to impact the amount of CPS produced (30, 45). To examine the role of carbon source on the CPS production, we quantified uronic acid, considered as a key component of *K. pneumoniae* CPS (**Fig. 5C**). As with mucoviscosity, *K. pneumoniae* produced high levels of CPS when grown in LB and reduced levels in MM irrespective of the carbon source, and addition of casamino acids to MM increased CPS. However, no differences were observed in either the amount of CPS or the transcriptional activation of the *cps* genes when comparing *K. pneumoniae* grown in MM supplemented with casamino acids with glucose or fucose as the carbon source (**Fig. 5C** **and S4A**). Thus, our data show that growth in MM with fucose as a carbon source positively modulates hypermucoviscosity without affecting capsule levels.

### Fucose utilization promotes auto-aggregation

*K. pneumoniae* is known to form aggregates as planktonic cells. This phenotype, known as auto-aggregation, is implicated in virulence in multiple pathogenic bacteria (46). To assess the ability of *K. pneumoniae* to auto-aggregate, we used a low-speed centrifugation assay distinct from that used to quantify HMV and calculating an Auto-aggregation Index (AI) based on the contribution of aggregated bacteria to optical density. We observed that growth in fucose resulted in enhanced auto-aggregation compared to growth in glucose or glycerol (**Fig. 6A**). Fimbriae are known contributors to auto-aggregation (46), however, the AI of a type 1 fimbriae mutant *(ΔfimH*) did not significantly differ from that of the WT when grown in either glucose or fucose (**Fig. S5**), suggesting that the well-characterized type 1 fimbriae does not significantly contribute to auto-aggregation. Next, we determined the contribution of type 3 fimbriae toward auto-aggregation. A reduced AI was observed with the type 3 fimbriae mutant (*ΔmrkABC*) irrespective of the carbon source (**Fig. S5**), but fucose-grown *K. pneumoniae* still displayed greater AI than glucose-grown *K. pneumoniae* (**Fig. 6B**). Furthermore, we did not observe a substantial increase in transcript abundance of either the type 1 or the type 3 fimbriae in fucose conditions (**Fig. S4B**). Taken together, our data suggest that the type 3 fimbriae are critical for planktonic cell-to-cell aggregation, though fucose-mediated aggregation does not occur through either the type 1 or the type 3 fimbriae.

**Fig 6.**
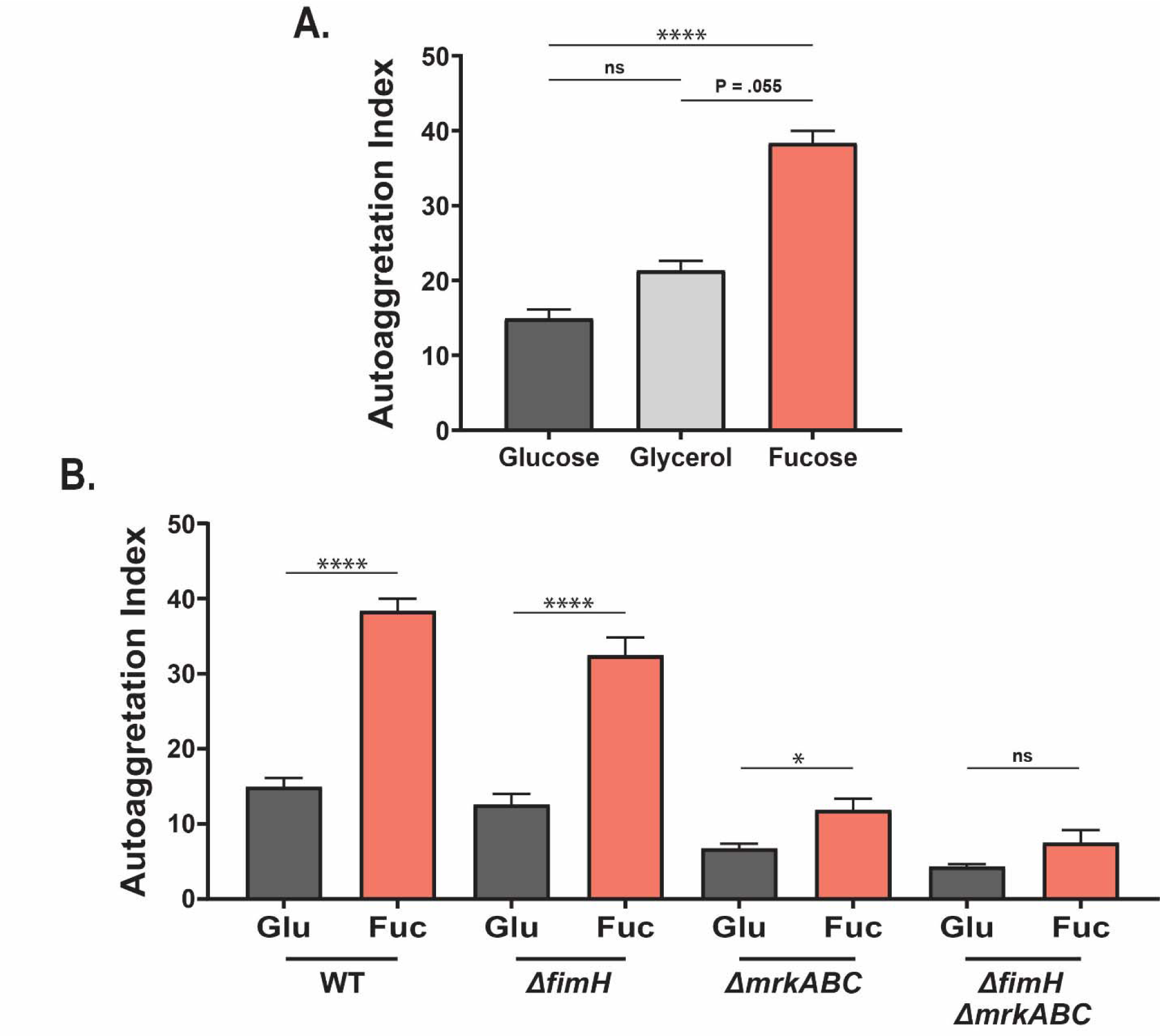
Fucose metabolism by *K. pneumoniae* enhances auto-aggregation. **(A)** Auto-aggregation of *K. pneumoniae* when grown in MM supplemented with either glucose, glycerol or fucose. **(B)** Contribution of type I and Type III fimbriae of *K. pneumoniae* towards auto-aggregation. WT, isogenic type I fimbriae mutant (*ΔfimH*), type III fimbriae mutant (*ΔmrkABC*), and the double fimbriae mutant (*ΔfimH, ΔmrkABC*) strains were tested in MM supplemented with either glucose or fucose. For each strain a Mann-Whitney *U* test was performed to compare glucose- and fucose-grown *K. pneumoniae*. For all graphs, shown is mean ± SEM from ≥3 individual experiments. * P < 0.05, **** P < 0.0001, ns - not significant

### Fucose utilization positively modulates biofilm formation

*K. pneumoniae* is known to readily form biofilms on solid surfaces including urethral catheters, which drives transmission and UTI incidence in the hospital setting (47). Additionally, enteric pathogens can form biofilms in the GI tract (48). Given the relevance of biofilms to colonization and nosocomial spread (49, 50), we used a microtiter dish colorimetric assay to investigate whether fucose utilization influences early biofilm formation. Compared to either glucose or glycerol, growth in fucose yielded increased biofilm formation (**Fig. 7A**). We further tested two unrelated clinical isolates to determine whether our results were strain-specific. Both isolates behaved similarly to KPPR1S and formed more robust biofilms when grown with fucose as the sole carbon source than with glucose (**Fig 7B**). Moreover, similar results were obtained with two *E. coli* isolates, revealing that the effect of fucose utilization on biofilm is conserved across multiple strains and species (**Fig. S6A**).

**Fig 7.**
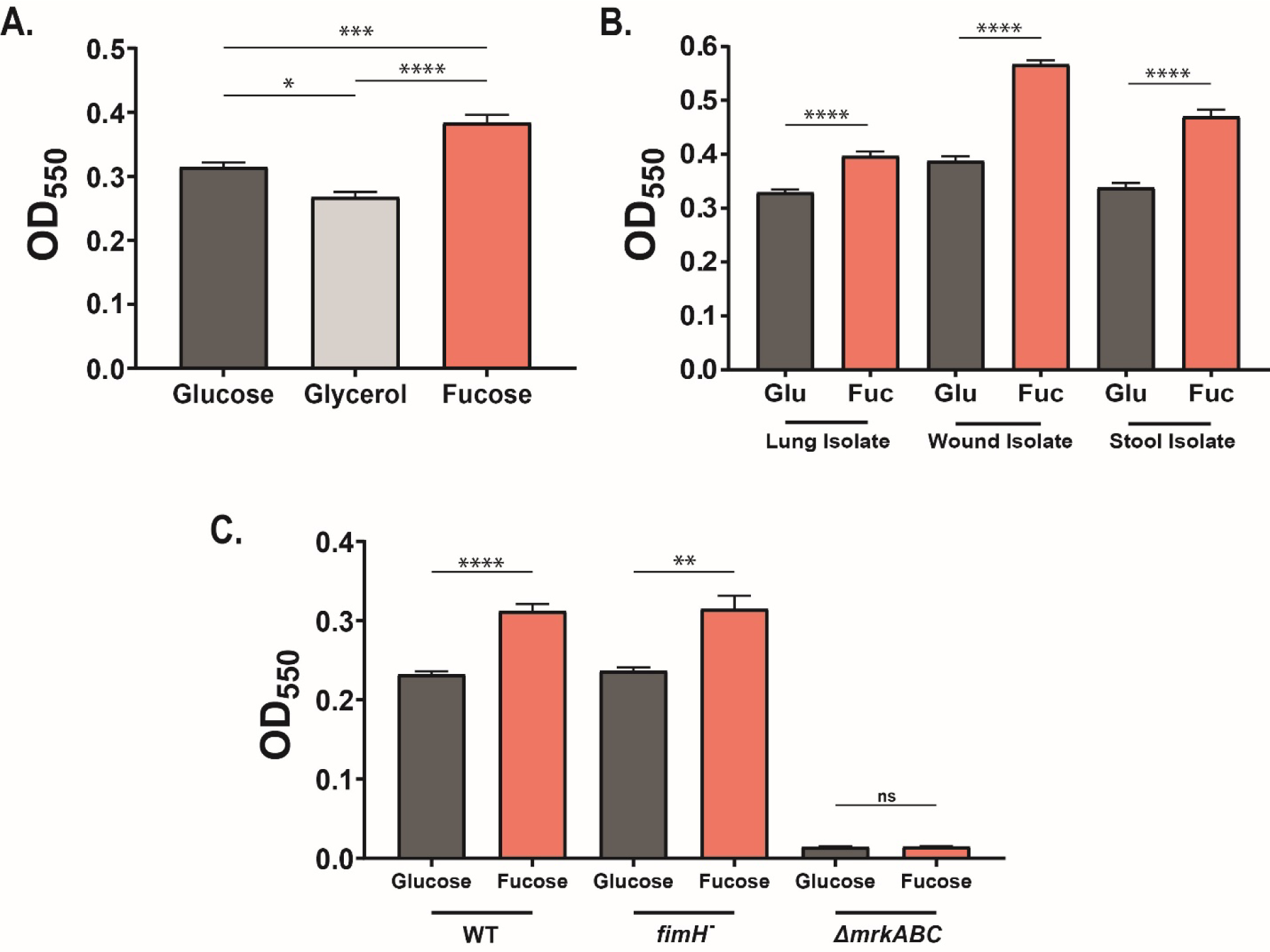
Fucose metabolism promotes biofilm formation in *K. pneumoniae*. **(A)** Comparison of *K. pneumoniae* (KPPR1S) biofilm produced when grown statically in MM supplemented with either glucose, glycerol, or fucose as the carbon source. To determine statistical differences a Kruskal-Wallis followed by Dunn’s test of multiple comparisons was performed. **(B)** Differences in biofilm amount produced by multiple clinical *K. pneumoniae* isolates. Biofilm grown in MM with either glucose or fucose as the carbon source was quantified for a clinical lung isolate (KPPR1S), a wound isolate (KP168), and a stool isolate (ST 37; *wzi* 108; K80). **(C)** Contribution of type I and type III fimbriae to biofilm production. Biofilm was quantified from fimbrial mutants, *ΔfimH* (type I fimbriae), and *ΔmrkABC* (type III fimbriae) of KPPR1S grown in MM with either glucose or fucose as the carbon source. A Mann-Whitney *U* test was performed to compare differences between glucose- and fucose-grown biofilms. Shown is mean ± SEM for ≥3 individual experiments. * P < 0.05, ** P < 0.01, *** P < 0.001, **** P < 0.0001, ns – not significant

Previous studies suggest that type 3 fimbriae are necessary for *K. pneumoniae* to form biofilms, while type 1 fimbriae are dispensable (51, 52). In agreement with these studies, we observed no change in biofilm production between the WT and the type 1 fimbrial mutant (*ΔfimH*). In contrast, irrespective of the carbon source, biofilm formation was entirely abrogated in the type 3 fimbrial mutant (*ΔmrkABC*), corroborating the importance of type 3 fimbriae in biofilm formation (52) (**Fig. 7C**).

### Enhanced fucose-mediated biofilms have increased proteinaceous EPS

The extracellular polymeric substances (EPS) of the biofilm matrix of *K. pneumoniae* consist of proteins, exopolysaccharides, and extracellular DNA (eDNA) (53, 54). Therefore, we sought to determine there were any key compositional differences in biofilms from cultures grown in glucose or fucose.

To provide insight into the composition of *K. pneumoniae* biofilms, we treated them with enzymes that targeted specific EPS components (54). Biofilms grown in glucose or fucose were treated with cellulase targeting cellulose, a known exopolysaccharide in *K. pneumoniae* biofilms (55); DNAse I, targeting eDNA; sodium metaperiodate (NaIO4), targeting β-1,6-N-acetyl-d-glucosamine (β-1,6-GlcNAc), another exopolysaccharide (56); and proteinase K, targeting proteins (**Fig. 8A-D**). A reduction in biofilm amount was observed only when treated with cellulase (**Fig. S6B**) and Proteinase K (**Fig. S6C**), suggesting cellulose and proteins constitute the major components of KPPR1S biofilm. However, the difference in biofilm robustness between glucose and fucose conditions remained unchanged after incubation with cellulase, suggesting that cellulose amount in biofilms grown in either glucose or fucose remains unchanged (**Fig. 8A**). Conversely, Proteinase K treatment led to abrogation of the difference observed between fucose- and glucose-grown biofilm, revealing that the increase in fucose grown biofilm results from higher protein content in the EPS (**Fig. 8D**). Collectively, our data indicate that fucose positively modulates biofilm formation by increasing the abundance of one or more matrix protein.

**Fig 8.**
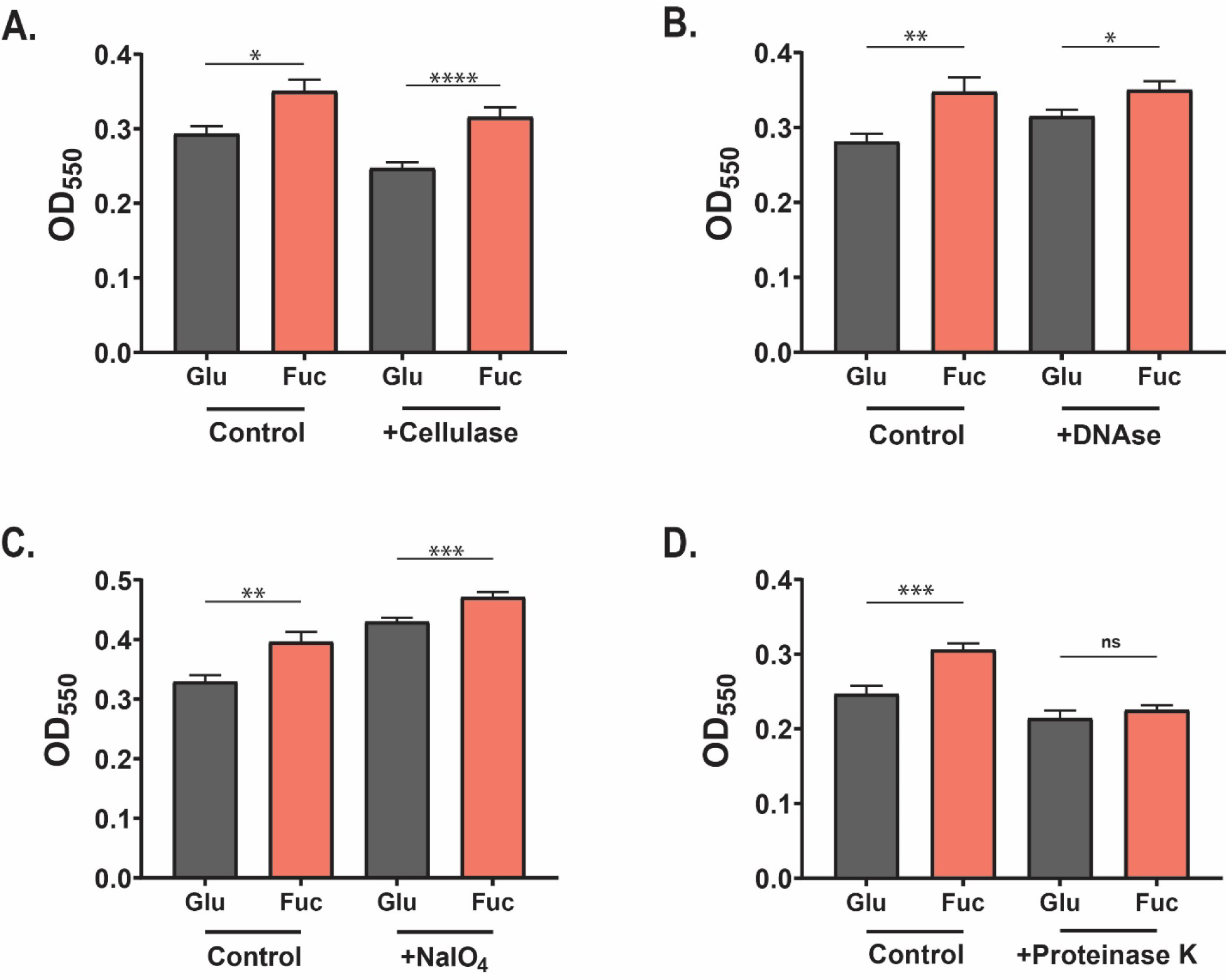
Fucose grown biofilms of *K. pneumoniae* have a higher proteinaceous component. **(A-D)** Enzymatic effect on biofilms to determine their biological content. Biofilms were grown in MM with either glucose or fucose as the carbon source, cells removed, and biofilms were treated with **(A)** Cellulase, **(B)** DNAse I, **(C)** Sodium metaperiodate (NaIO_4_), and **(D)** Proteinase K, or with their corresponding buffers (control), and biofilms quantified. See Materials and Methods for details. A Mann-Whitney *U* test was performed comparing glucose- and fucose-grown biofilms. For all graphs, shown is mean ± SEM. * P < 0.05, ** P < 0.01, *** P < 0.001, **** P < 0.0001, ns – not significant

## Discussion

The focus of the current study was on the ability of *Klebsiella pneumoniae* to metabolize alternative carbon sources present in the gut to evade CR and modulate its virulence phenotypes. We focused on fucose utilization by *K. pneumoniae* for two reasons. Firstly, fucose is abundant in the GI tract, with the majority of fucose being mucin-derived (57). Secondly, fucose can modulate the expression of virulence genes in gut pathogens (27). Fucosidase-producing gut commensals such as *B. thetaiotaomicron* liberate fucose to metabolize the sugar and incorporate it into its capsular polysaccharide (58), but unused fucose remains free to be utilized by other microbes unable to cleave fucose from mucin, such as *K. pneumoniae* (20-22). Using our murine model of *K. pneumoniae* gut colonization with an intact microbiota allowed us to further our understanding of how a silent gut colonizer metabolizes liberated fucose to bypass CR. Both in single and co-infection GI colonization studies, the WT strain out-competed and shed more robustly than the strain defective in fucose metabolism (*ΔfucI*). When the *ΔfucI* genotype was chromosomally restored (*fucI^+^*), it behaved as WT, suggesting the defect in gut colonization was specific to fucose metabolism.

In a hospital setting, patients colonized with *K. pneumoniae* tend to be on antibiotics (6, 59). Antibiotic treatment reduces bacterial diversity and density in the GI tract, triggering an increase in concentration of mucin-derived polysaccharides in the murine gut as consumption by native flora is abrogated (22). Thus, we examined whether fucose utilization by *K. pneumoniae* was dispensable in the GI tract during antibiotic pressure. *S. typhimurium*, a prototypic murine gut pathogen, metabolizes mucin-derived fucose and other glycans to colonize the GI tract in conventional mice treated with antibiotics (22). However, unlike *S. typhimurium*, *K. pneumoniae* does not require fucose utilization for post-antibiotic expansion, as both the WT and *ΔfucI* strains expand to equally high density (supershedder) in the antibiotic-treated gut (**Fig. 3D**). It is likely that *K. pneumoniae* relies on different carbon sources when nutritional competition is reduced, suggesting that fucose utilization assists in avoiding CR by providing an alternative carbon niche underutilized by other members of the microbiota. Our results are supported by a recent transposon mutagenesis screen that suggest that the fucose operon of *K. pneumoniae* is dispensable for gut colonization during antibiotic pressure (60).

In addition, our growth results in CF suggest a potential role for probiotics in preventing *K. pneumoniae* colonization. Recently, *K. oxytoca* was demonstrated to provide CR against a *K. pneumoniae* MDR isolate (15). EcN has previously been shown to protect the murine GI tract from EHEC colonization (3, 61), and at 24 hours post-inoculation likewise out-competed both the WT and *ΔfucI* strains when grown *in vitro*. This out-competition was observed in the physiologically relevant environment of CF, but not in MM supplemented with glucose or fucose, indicating that the ability to provide CR depends greatly on the metabolic landscape. CF supplies a complex nutritional milieu (15), and the observation that EcN prevails in this media suggests that it utilizes certain nutrients present in the GI tract more efficiently than *K. pneumoniae.* Future studies will aim to define the underlying molecular mechanism of EcN-dependent out-competition of *K. pneumoniae*.

Pathogens often use the carbohydrate landscape in the gut to contextualize their position within a host and control virulence accordingly (27, 62). Therefore, we examined whether fucose metabolism modulated the virulence phenotypes of *K. pneumoniae* and observed that it enhanced the HMV phenotype, considered a hallmark of hypervirulent *K. pneumoniae* strains. Walker *et. al* have shown that although CPS production is required for HMV, the two phenotypes are genetically distinct (43). Recently, a deletion of *mgrB* in *K. pneumoniae* affected capsule amount, but did not impact HMV (44). Our results herein further validate that the two phenotypes are separate. Increase in HMV in fucose conditions was associated with an upregulation of the *rmp* locus, which positively modulates capsule (*rmpC*) and HMV (*rmpD*) production (41, 43). Our study is the first to show that carbohydrate utilization can affect HMV without impacting CPS, providing further insight into the complex regulation of the *rmp* locus. We did observe *rmpC* to be upregulated in fucose growth conditions, without a corresponding increase in CPS levels. However, a lack of corresponding increase in CPS can be because the *rmpC* transcript is potentially post-transcriptionally regulated, or our CPS quantification assay is not sensitive enough to detect subtle changes in CPS amount.

To provide a molecular insight into how fucose modulates biofilms, we determined the structural content of biofilm through enzymatic assays and showed that the difference observed between glucose- and fucose-grown biofilms could be attributed to an increase in matrix protein content. These may include eDNA binding proteins or other fimbriae. Besides type 1 and 3 fimbriae, the *K. pneumoniae* genome contains multiple loci encoding other fimbrial structures including *E. coli* Common Pilus (63-66), which could contribute towards increased biofilm and auto-aggregation under fucose growth conditions.

Gut colonization is the first step in the pathogenesis and host-to-host transmission of *K. pneumoniae.* Thus, identifying the strategies that *K. pneumoniae* employs for colonization is crucial to combating this pathogen. The present study provides a molecular understanding of the role of the alternative carbon source fucose on gut colonization and virulence of *K. pneumoniae*. We show that fucose liberated by gut commensals serves as an important energy source for *K. pneumoniae*, allowing it to bypass nutritional competition in the gut. Furthermore, we demonstrate that EcN can potentially be used to enhance CR against *K. pneumoniae*, as they likely occupy the same metabolic niche. Lastly, we provide insights into the complex relationship between carbon metabolism and the modulation of virulence, which could provide a framework for strategies to combat the spread of this pathogen.

## Materials and Methods

### Ethics Statement

This study was conducted according to the guidelines outlined by National Science Foundation animal welfare requirements and the Public Health Service Policy on Humane Care and Use of Laboratory Animals (67). The Wake Forest Baptist Medical Center IACUC oversees the welfare, well-being, and proper care and use of all vertebrate animals. The approved protocol number for this project is A20-084.

### Bacterial strain construction

Strains, plasmids and primers used in this study are listed in **Table 1** and **Tables S1-2**, respectively. To construct an in-frame gene deletion of *fucI* (*fucI::cam*) in KPPR1S background (34), genomic DNA was isolated from the *fucI*::cam mutant in the MKP103 background (68), and PCR was performed using Q5 polymerase (NEB; M0491L) with primers that had 500 bp homology to both KPPR1S and MKP103 on either end of the transposon cassette. The purified PCR product was then electroporated into the target strain (AZ63, **Table 1**) containing the temperature sensitive plasmid pKD46 (**Table S1**), which contained λ red recombination genes downstream of an arabinose inducible promoter. Recombination was carried out as described (69) to generate *fucI*::cam KPPR1S (AZ120). Mutants were isolated on agar plates containing chloramphenicol (50 *µ*g/ml), and single colonies were purified and verified through PCR. Removal of the chloramphenicol cassette was done as previously described using the plasmid pCre2 (68) to generate a clean deletion (AZ170) used as *ΔfucI* except where otherwise stated. To construct the *fimH*::cam, *ΔmrkABC* double mutant (AZ203), genomic DNA from a *fimH*::cam mutant (AZ108) was used as Q5 PCR template and λ red recombination was performed as described above. The resultant mutants were purified and subsequently verified through PCR. The chromosomal complement of *ΔfucI* (*fucI^+^*, AZ171) was constructed as previously described (69) with slight modification. Briefly, the *fucI* gene and ∼500 bp upstream and downstream was amplified using WT (AZ55) genomic DNA as a template with primers than contain homology sequences to the plasmid pKAS46 (**Table S1**). The plasmid was then digested with NotI and NheI, and the *fucI* PCR product was cloned into the plasmid using the NEBuilder HiFi DNA Assembly Kit (NEB; E5520S), followed by transformation into the *E. coli* strain S17-1 *λpir*. Conjugation with the *ΔfucI* strain and subsequent selection of the complemented strain were carried out as previously described (70).

**Table 1.**
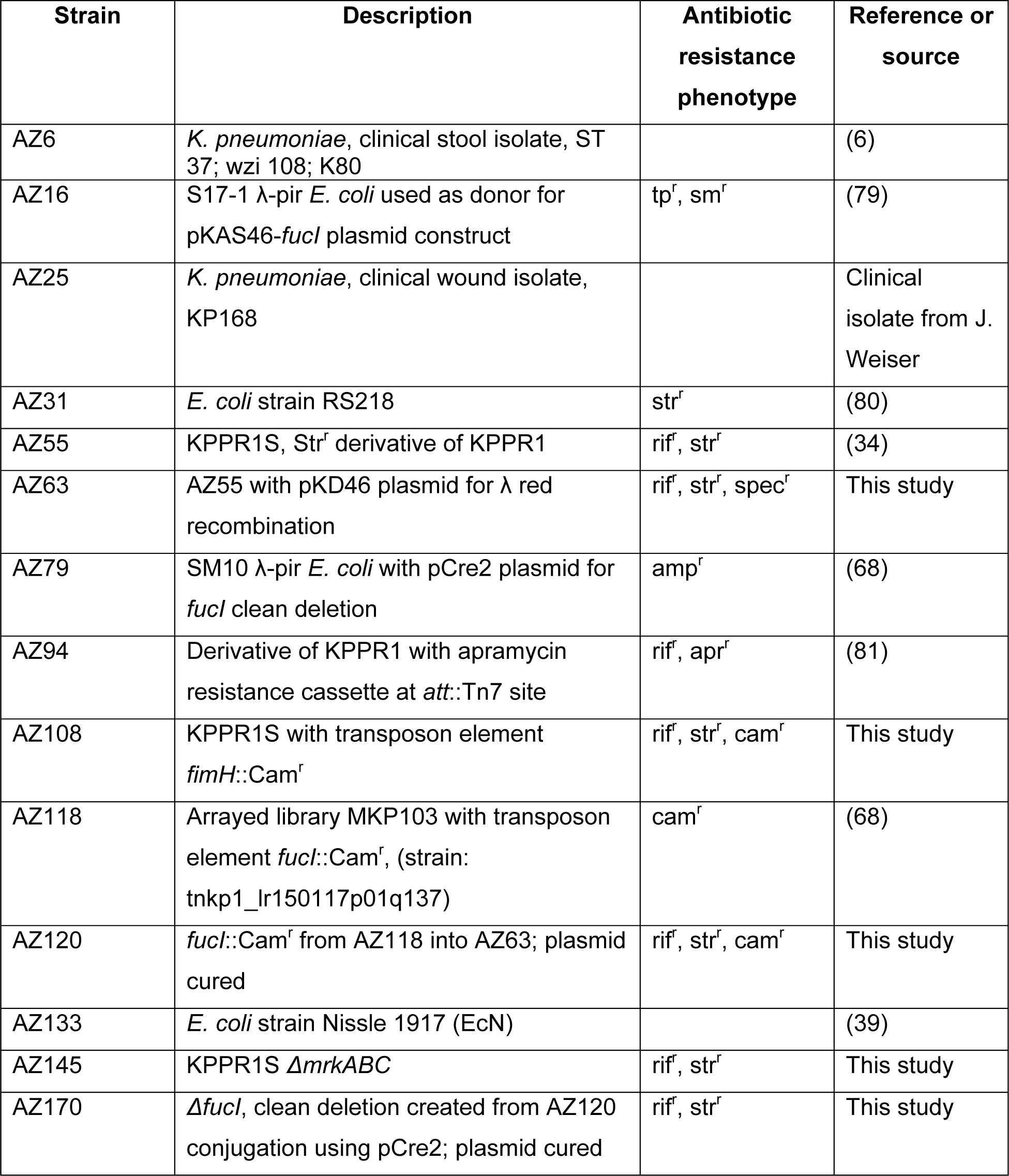
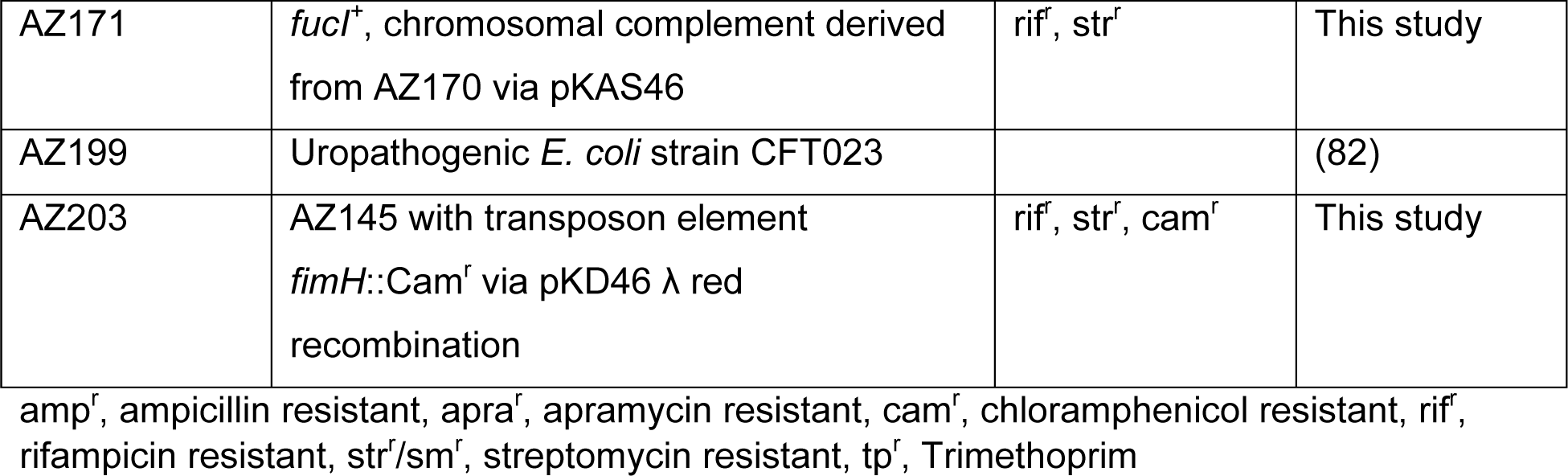
Strains used in the study

### Preparation of Cecal Filtrate (CF)

To prepare CF, cecal contents from euthanized mice were placed in screw-cap tubes (Fisherbrand; 02-682-558) with 2-3 glass beads (BioSpec Products; 11079127), diluted 1:5 (weight to volume) in PBS, homogenized using a beadmill, and centrifuged (20,000 *x g* for 10 min). The supernatant was then transferred to a separate tube and centrifuged again. The resulting supernatant was passed through a 0.2 *µ*m nylon syringe filter (Fisherbrand; 09-719C). Prior to inoculation the CF was further diluted with PBS to a final concentration of 1:10 weight to volume.

### Growth curves, generation time, and Competition Experiments

For growth studies, KPPR1S was grown overnight at 37°C with constant agitation in LB-Lennox, or in M63 minimal media (MM) supplemented with either 0.5% glucose, glycerol, or fucose as sole carbon source. Cultures were diluted 1:100 into their respective fresh growth media and grown at 37°C with constant agitation. For growth studies in CF, overnight cultures in LB were spun down and washed with 1X PBS, diluted 1:100 into CF and grown likewise. At designated time-points, a sample was removed, diluted, and plated for enumeration.

To calculate generation time in LB and MM, strains were grown as above until the samples reached an OD_600_ of 0.2, following which a sample was removed, diluted, and plated for enumeration every 15 minutes for up to an hour. The generation time was calculated (44, 71) using the formula G=t/n, where G is the generation time, t is the time interval, and n is the number of generations as calculated by the formula:

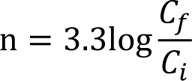

Where C_f_ is the quantity of bacteria at the end of a time interval and C_i_ is the amount at the beginning of the time interval.

For competition experiments in MM, overnight cultures of competing strains were diluted 1:100 each into a single culture of MM supplemented with glucose and grown at 37°C with constant agitation. At inoculation and at designated time-points, a sample was removed, diluted, and plated on selective antibiotics for enumeration of each strain. The Competitive Index (CI) for two strains was calculated at each time-point using the following formula (72):

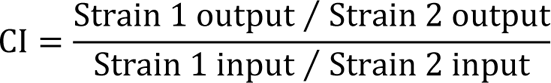

For competition experiments in CF, strains were grown overnight in LB, spun down and washed with 1X PBS, then diluted 1:100 each into fresh CF In competition experiments between *K. pneumoniae* and *E. coli* strain Nissle 1917, colonies were easily distinguished by size and morphology in lieu of antibiotic selection. CI was calculated as described.

### Mouse inoculations for colonization, shedding, and qRT-PCR

C57BL/6J Specific Pathogen Free mice (SPF) were obtained from Jackson Laboratory (Bar Harbor, ME), bred and maintained in the animal facility at Biotech Place, Wake Forest Baptist Medical Center. Mice were inoculated as described (12). Briefly, 5-7 week old mice had food and water removed for 4 hours, and were then fed *K. pneumoniae* in 2% sucrose-PBS via pipette tip in two 50-*µ*l doses at ∼10^6^ CFU/100 *µ*l. The inoculation dose was serially diluted and plated on selective antibiotic plates to determine the exact dose. The mice were monitored through the time course of the experiment.

Fecal collection was carried out as described (12). To enumerate the bacterial shedding, fecal pellets were weighed and placed in screw-cap tubes (Fisherbrand; 02-682-558) with 2-3 glass beads (BioSpec Products; 11079127), diluted 1:10 (weight to volume) in 1X PBS, homogenized using a beadmill, serially diluted, and plated on selective antibiotic plates. The limit of detection for fecal shedding was 10^2^ CFU/ml.

To induce supershedder phenotype mice were inoculated as described (12), and 5 days post-inoculation a single dose of streptomycin (in sterile PBS) was administered via oral gavage (5 mg/200 *µ*l). *K. pneumoniae* was enumerated from fecal shedding for an additional 10 days.

To determine colonization density in the GI tract, the cecum, ileum, and colon were removed from mice following CO_2_ (2 L/min for 5min) euthanasia with subsequent cardiac puncture. Sections of the proximal colon and terminal ileum as well as the whole cecum were removed from each mouse. As described above, GI samples were weighed, placed in screw-cap tubes with beads, diluted 1:10 (weight to volume) in PBS, homogenized, serially diluted, and plated. The limit of detection for organ homogenate was 10^2^ CFU/ml.

Oropharyngeal lavage was carried out with 200 *μ*l of sterile PBS. The esophagus was exposed and cut transversely. A gavage needle, attached to a prefilled insulin syringe (BD) with 1× PBS, was then inserted into the cut esophagus, and PBS was collected from the mouth. The collected lavage was serially diluted and plated on appropriate antibiotic plates. The limit of detection for oral lavage was 33 CFU/ml.

Inoculation and fecal shedding collection for competition studies (Competition Index [CI]) were conducted as described above with the inoculation dose containing a 1:1 ratio of *ΔfucI K. pneumoniae* (AZ170) and an apramycin resistant derivative of KPPR1 (AZ94). The CI was calculated using the following formula:

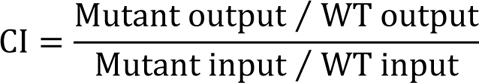

### RNA extraction, cDNA synthesis, and quantitative reverse transcription PCR

RNA was isolated from *K. pneumoniae* grown *in vitro* using the trizol method as described (73) with slight modifications. Overnight cultures grown in MM + 0.5% glucose or fucose were diluted into their respective media and grown to OD_600_ ∼0.5; for *rmp* expression analysis, 0.2% casamino acid was added to the overnight and subsequent cultures.

For bacterial RNA isolation from the gastrointestinal tract of mice, a modified protocol (74) was used. Cecal contents were removed from colonized mice and placed into a 2 ml screwcap tube (Fisherbrand; 02-682-558) containing two glass beads (2.7 mm; BioSpec Products; 11079127). RNAlater (Invitrogen; AM7020) was then added at a volume of 1 ml / 1 g of cecal contents. After homogenization, samples were incubated overnight at 4°C. A volume of chilled 1X PBS was added equal to the volume of RNALater, samples were centrifuged (700 *x g* for 1 min at 4°C), and the supernatant was transferred to a new tube and centrifuged again (900 *x g* for 5 min at 4°C). The supernatant was then discarded, and the pellet was resuspended in 500 *µ*l 2x Buffer A (200 mM NaCl, 200 mM Tris, 20 mM EDTA), 210 *µ*l 20% SDS, and 500 *µ*l acid phenol:chloroform:isoamyl alcohol (25:24:1) solution (Invitrogen; 15593-031), added to screwcap tubes containing 250 *µ*l 0.1-mm-diameter silica beads (BioSpec Products) and the sample homogenized (1 min, 4.5 strength). Afterwards, samples were centrifuged (6,000 *x g* for 3 min at 4°C), and the aqueous layer was transferred to a 2 ml phase lock gel tube (QuantaBio) alongside an equal volume of phenol:chloroform:isoamyl alcohol (25:24:1) solution (Invitrogen; 15593-031). Samples were centrifuged (18,000 *x g* for 5 min at 4°C), the aqueous phase was transferred to a new tube alongside 600 *µ*l isopropanol, 60 *µ*l of 3M sodium acetate, and 5 *µ*g glycogen, and incubated overnight at -20°C. Afterwards samples were centrifuged (18,000 *x g* for 20 min at 4°C), and the resultant pellet washed three times between with 500 *µ*l 100% ethanol, dried, and subsequently resuspended in RNase free H_2_O.

Total RNA from broth cultures and cecal samples were treated with DNase (Invitrogen; AM1907) per the manufacturer’s instructions and cDNA was synthesized as described (73). Quantitative reverse transcription PCR (qRT-PCR) was conducted as previously described (75, 76) with slight modifications. iTaq Universal SYBR Green Supermix (Bio-Rad; 1725121) was used with 20 ng cDNA and 0.5 mM primers per reaction. Samples were run in duplicate on a CFX384 Touch real-time PCR detection system (Bio-Rad). For *in vitro* experiments, primers directed toward *gyrA* were used as the internal control. Comparison of *in vivo* (cecal) to *in vitro* (M63 + 0.5% glucose) fold-change gene expression was performed with *K. pneumoniae*-specific primers targeting the 16S subunit for internal control. RNA expression was quantified using the ΔΔC_T_ threshold cycle (C_T_) method (75). Fold-change was calculated as 2^ΔΔC^T. For *in vivo* gene expression, fold-change values were further normalized to *gyrA* expression to reduce variability.

### Mucoviscosity assay

Mucoviscosity was determined using the low-speed centrifugation assay previously described (41). Briefly, cultures were grown in the indicated media as described above, and the OD_600_ adjusted to 1. Samples were then centrifuged (1000 *x g* for 5 min), and the OD_600_ of the supernatant was measured.

### Uronic acid assay

Uronic acid content as an indicator of capsular polysaccharide amount was quantified as described previously (41). Cultures were grown in indicated media, and uronic acid extracted by adding 100 *μ*l 1% zwittergent + ∼25% citric acid to 500 *μ*l culture and the mixture incubated at 50°C for 20 min. Samples were centrifuged (20,000 *x g* for 5 min) and 300 *μ*l supernatant was incubated overnight with 1.2 ml 100% ethanol. Precipitated material was then pelleted by centrifugation (20,000 *x g* for 5 min), resuspended in 200 *μ*l H_2_O, to which 1.2 ml sulfuric acid + 12.5 mM sodium tetraborate solution was added and the sample chilled on ice for 10 min. Afterwards, 20 *μ*l 2-phenylphenol was added to the samples that acts as colorimetric indicator uronic acid presence. The OD_520_ was measured and analyzed with a standard curve generated by glucuronic acid.

### Auto-aggregation assay

Auto-aggregation was quantified by a low-speed centrifugation assay (77) with modifications. *K. pneumoniae* was grown in MM supplemented with either 0.5% glucose, glycerol, or fucose as the carbon source at 37°C with constant agitation for 30 hours. Cultures were gently vortexed for even distribution of bacterial aggregates (flocs), and 1 ml immediately transferred into a 1.5 ml Eppendorf tube. The tube was gently centrifuged for five seconds on a benchtop microfuge to pellet the flocs. 100 *μ*l supernatant was removed and its OD_600_ measured (Supernatant OD_600_). The 30-hour culture containing flocs was then vortexed to disrupt and homogenize the flocs, and OD_600_ was measured (Vortexed OD_600_). An auto-aggregation index was calculated using the following formula:

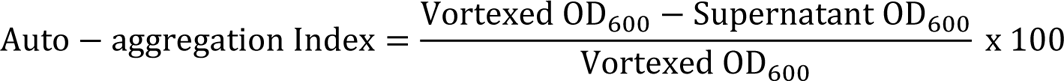

### Biofilm assay

Quantification of mature biofilm was done as described (78) with adjustments. Briefly, cultures were grown in fresh LB-Lennox overnight and diluted 1:100 in indicated media. 100 *μ*l of the bacterial suspension was placed in the wells of a 96-well round-bottom culture plate (Costar; 3799), sealed with thermal adhesive sealing film (Fisherbrand; 08-408-240), and incubated for 24 hours at 37°C. Afterwards, the plate was inverted and gently shaken to remove planktonic bacteria, washed twice by submerging the plate in a tub of diH_2_O and blotted on absorbent pads to remove excess liquid. Staining and quantification were carried out as described (78). Enzymatic treatment of mature biofilms was performed after removal of planktonic bacteria. Wells were treated with either 100 *μ*l of buffer with the specified enzyme or buffer alone, and the plate was incubated at 37°C. Plates were incubated for 1 hour with 100 *μ*g/ml proteinase K (Thermo Scientific) in pH 5.0 sodium acetate buffer, 1% cellulase (MP Biomedicals; 150583) in pH 5.0 sodium acetate buffer, or 40 mM sodium metaperiodate (NaIO_4_) (Fisher Chemical; S398-50) in pH 4.5 sodium acetate buffer. For DNAse treatment, plates were incubated for 24 hours with 100 *μ*g/ml DNase I (Alfa Aesar; J62229-MB) in PBS. After enzymatic treatment, the plates were washed, stained and quantified as described above.

### Statistical analysis

All statistical analyses were performed using GraphPad Prism (version 9.0) software (GraphPad Software, Inc., San Diego, CA). Unless otherwise specified, differences were determined using the Mann-Whitney *U* test (comparing two groups) or the Kruskal-Wallis test with Dunn’s test of multiple comparisons.

## Acknowledgements

We would like to thank Melissa Kendall (UVA School of Medicine) for providing the *E. coli* Nissle isolate. We would also like to thank Virginia Miller (UNC Chapel Hill) for providing the strain KPPR1S *ΔmrkABC* and for fruitful discussions pertaining to this manuscript, and Kimberly Walker (UNC Chapel Hill) for valuable conversations and input on the project. This project was supported by startup funds and graduate program funds provided by Wake Forest Baptist Medical Center to M.A.Z and A.W.H. A.S.B was supported by NIAID training program in immunology and pathogenesis (5T32AI007401-28).

---

**Fig S1.**
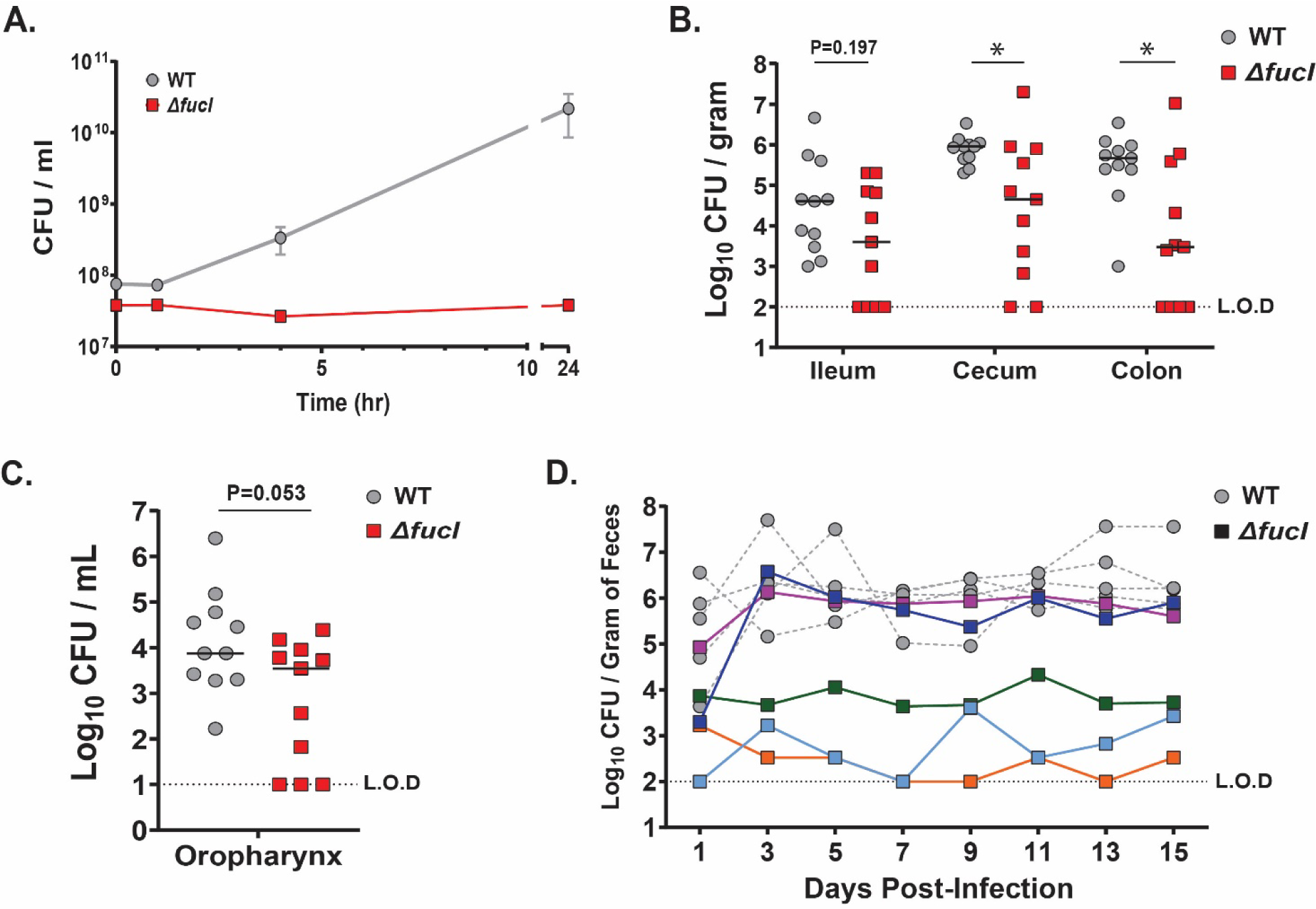
**(A)** Growth dynamics of the WT and isogenic *ΔfucI* mutant in MM supplemented with fucose represented as CFU/ml as a function of time. Mean ± SEM for ≥3 independent experiments is shown. **(B-C)** Colonization density of the WT and the *ΔfucI* strain day 15 post-infection in **(B)** ileum, cecum, colon, and **(C)** oropharynx. A Mann-Whitney *U* test was performed to compare colonization density between strains. Bars indicate median colonization density, and the dashed line indicates limit of detection (L.O.D). **(D)** Shedding dynamics of individual mice colonized with either the WT or the *ΔfucI* strain. Fecal shedding from select mice colonized with either the WT or *ΔfucI* strain shown in Fig. 3A with lines connecting individual mice. Each symbol represents a single mouse on a given day and the dashed line indicates limit of detection (L.O.D). * P < 0.05

**Fig S2.**
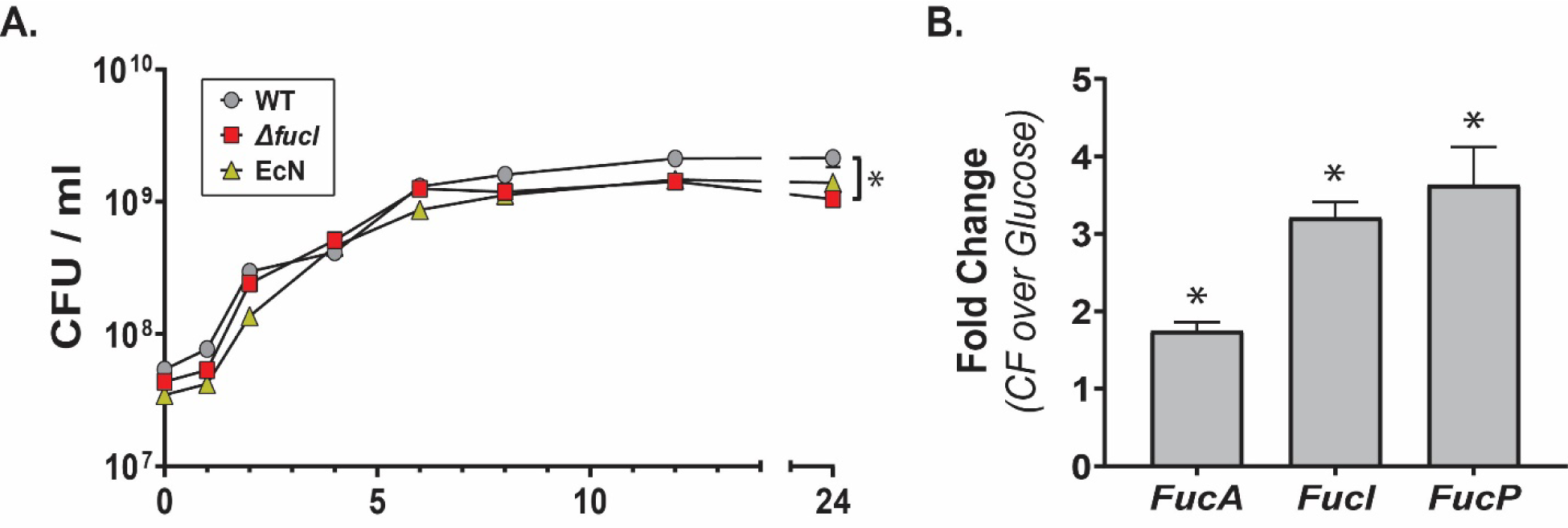
**(A)** Growth kinetics of the *K. pneumoniae* WT, *ΔfucI*, and EcN when grown individually in CF. Mean ± SEM for ≥3 independent experiments is shown. A Mann-Whitney *U* test was performed to compare between the WT and *ΔfucI* mutant at 24 hours. **(B)** qRT-PCR comparing *fuc* operon gene expression from *K. pneumoniae* grown in either CF or MM with glucose as the carbon source. Shown is fold change. *gyrA* was used as the housekeeping gene for 2*^-ΔΔC^T* analysis. For all graphs, mean ± SEM for ≥3 independent experiments is shown. * P < 0.05

**Fig S3.**
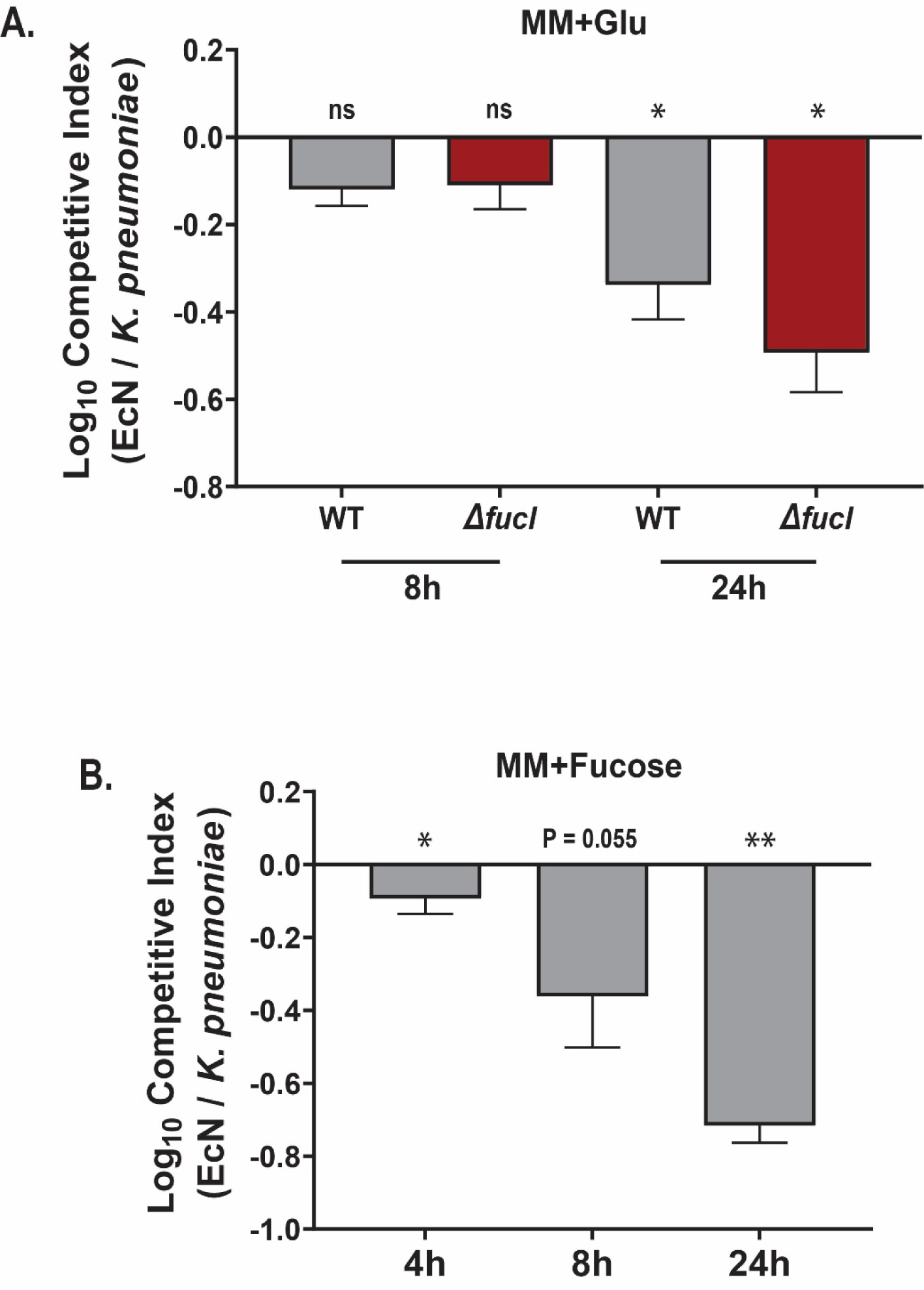
Competition experiments between *K. pneumoniae* and EcN in MM. Either *K. pneumoniae* WT or the *ΔfucI* strain were inoculated with EcN at a 1:1 ratio into MM supplemented with **(A)** glucose or **(B)** fucose, with CFUs enumerated and CI values calculated **(A)** at 8 and 24 hours post-inoculation or **(B)** 4, 8 and 24 hours post-inoculation. Shown is mean ± SEM from ≥3 individual experiments. Statistical significances of CIs were calculated using Wilcoxon signed-rank test with a theoretical median of 0. * P < 0.05, ** P < 0.01, ns - not significant

**Fig S4.**
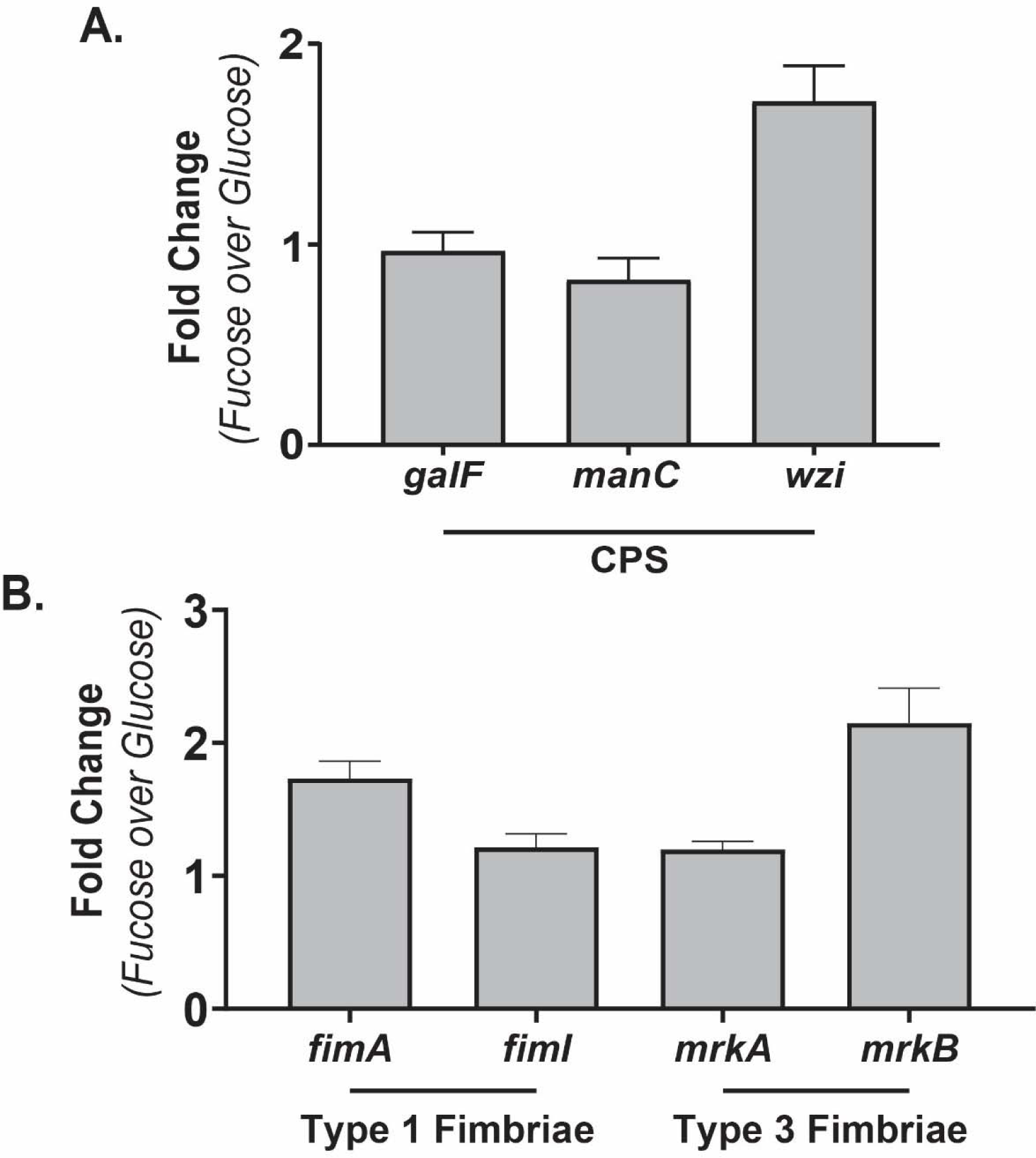
qRT-PCR comparing **(A)** CPS locus and **(B)** fimbriae locus gene expression from *K. pneumoniae* grown in MM with either glucose or fucose as the carbon source. Shown is fold change difference mean ± SEM. *gyrA* was used as the housekeeping gene for 2*^-ΔΔC^T* analysis.

**Fig S5.**
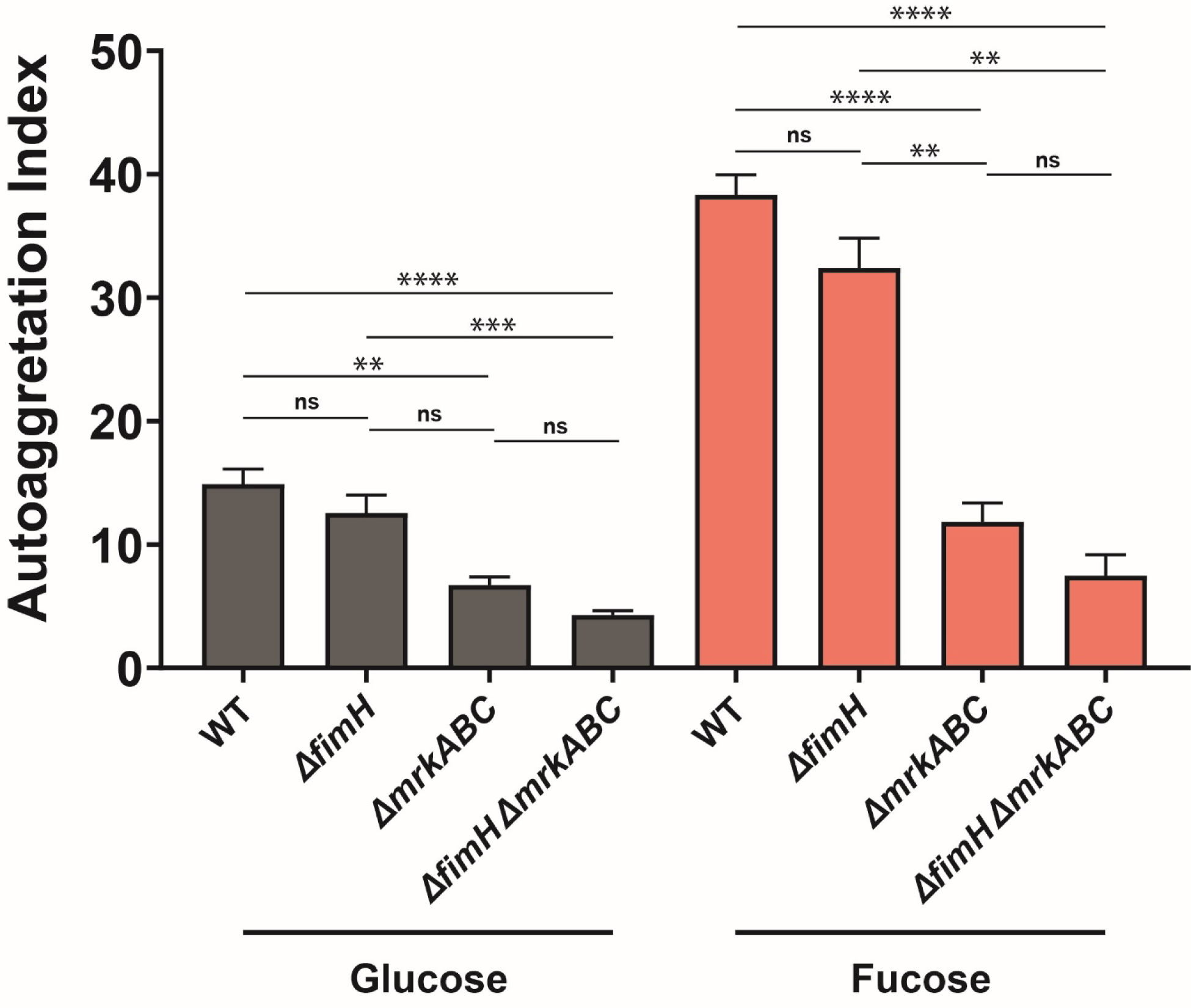
Evaluation of auto-aggregation data from Fig. 6B comparing AI of WT and fimbriae mutants when grown in specific carbon source. For each carbon source a Kruskal-Wallis followed by Dunn’s test of multiple comparisons was performed to compare between strains. For all graphs, shown is mean ± SEM from ≥3 individual experiments. ** P < 0.01, *** P < 0.001, **** P < 0.0001, *ns* – not significant

**Fig S6.**
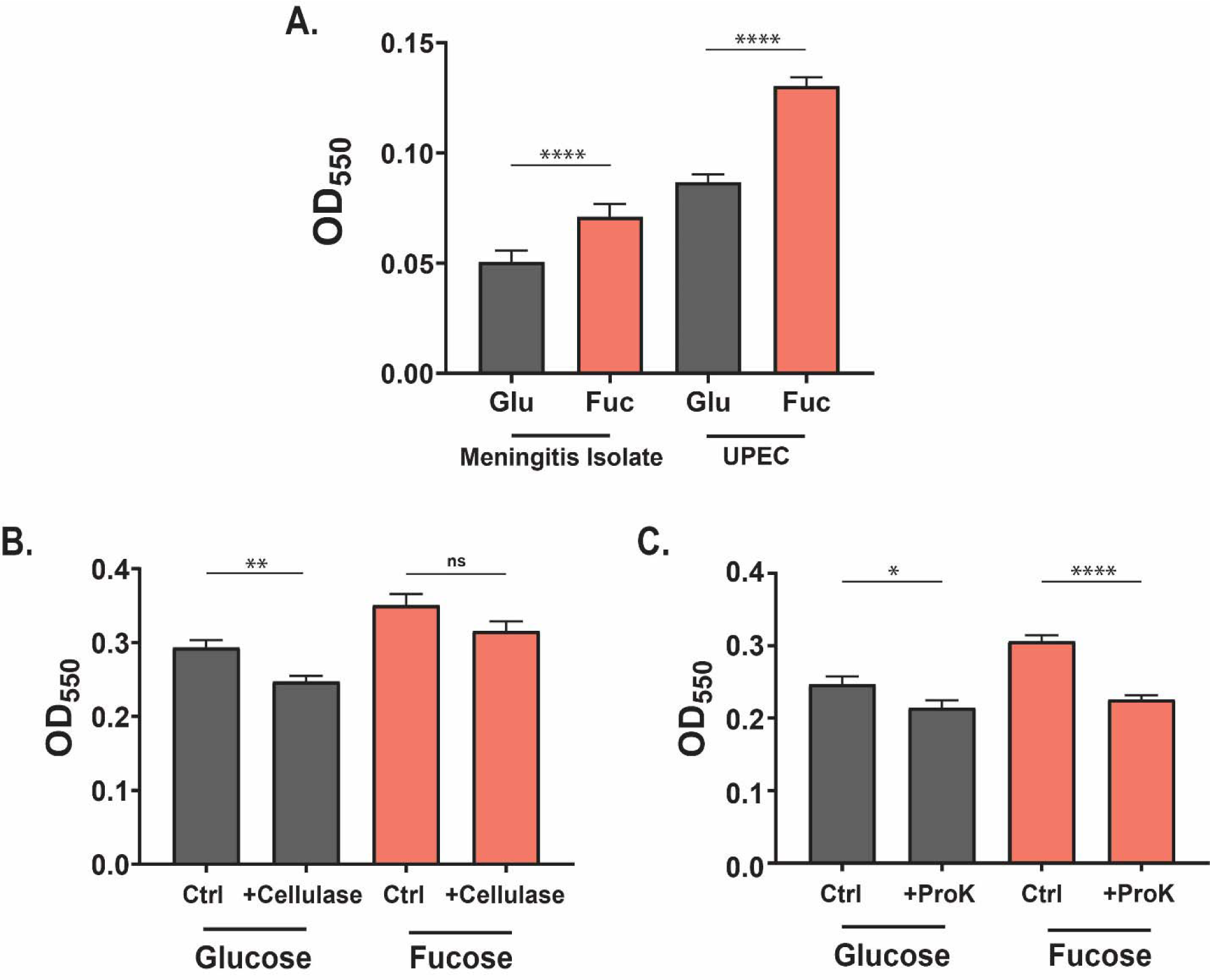
**(A)** Differences in biofilm amount produced by two clinical *E. coli* isolates. A neonatal meningitis isolate (RS218) and a uropathogenic *E. coli* (UPEC) isolate (CFT023) isolates were grown statically for 24 hrs in MM with either glucose or fucose as the carbon source and biofilm quantified. For each strain a Mann-Whitney *U* test was performed to compare between glucose- and fucose-grown *E. coli*. **(B)** The effect of Cellulase and **(C)** Proteinase K (ProK) on *K. pneumoniae* grown biofilms, as shown in Fig. 8A**-D**. A Mann-Whitney *U* test was performed comparing enzyme-treated and untreated biofilms. For all graphs, shown is mean ± SEM for ≥3 individual experiments. * P < 0.05, ** P < 0.01, **** P < 0.0001, ns – not significant

**Table S1.** Plasmids used in the study.

| Plasmid | Description | Antibiotic resistance phenotype | Reference or source |
| --- | --- | --- | --- |
| pKD46 | $\lambda$ red recombinase genes downstream of *araBAD* promoter; temperature sensitive (32°C) | spec<sup>r</sup> | (69) |
| pKAS46 | Vector for allelic exchange, contains *rpsL* for streptomycin counter-selection | kan<sup>r</sup>, str<sup>s</sup> | (70) |
| pCre2 | Cre recombinase for removing loxP *cam* cassette | amp<sup>r</sup> | (68) |

amp<sup>r</sup>, ampicillin resistant, kan<sup>r</sup>, kanamycin resistant, spec<sup>r</sup>, spectinomycin resistant, str<sup>s</sup> streptomycin sensitive

**Table S2.**
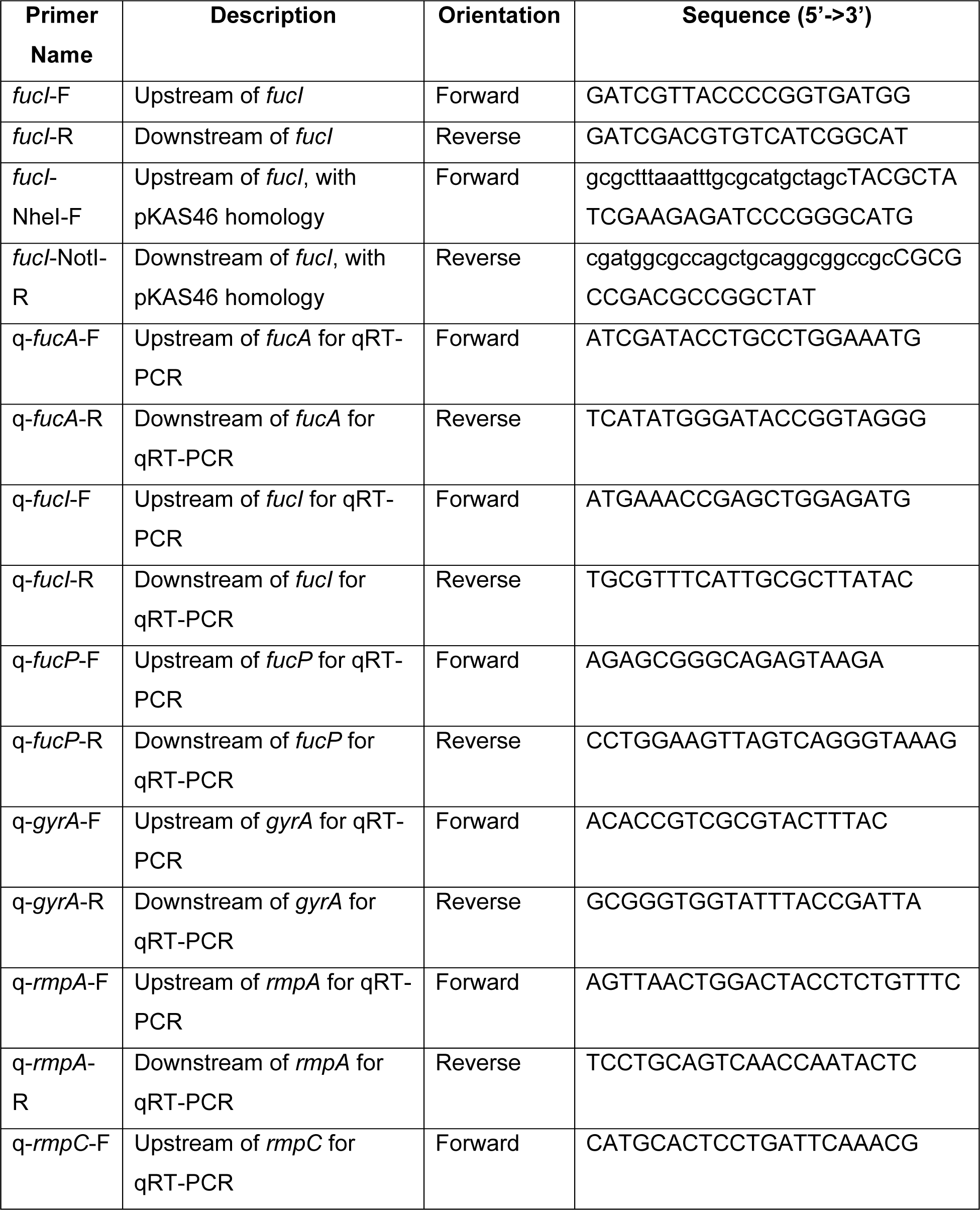
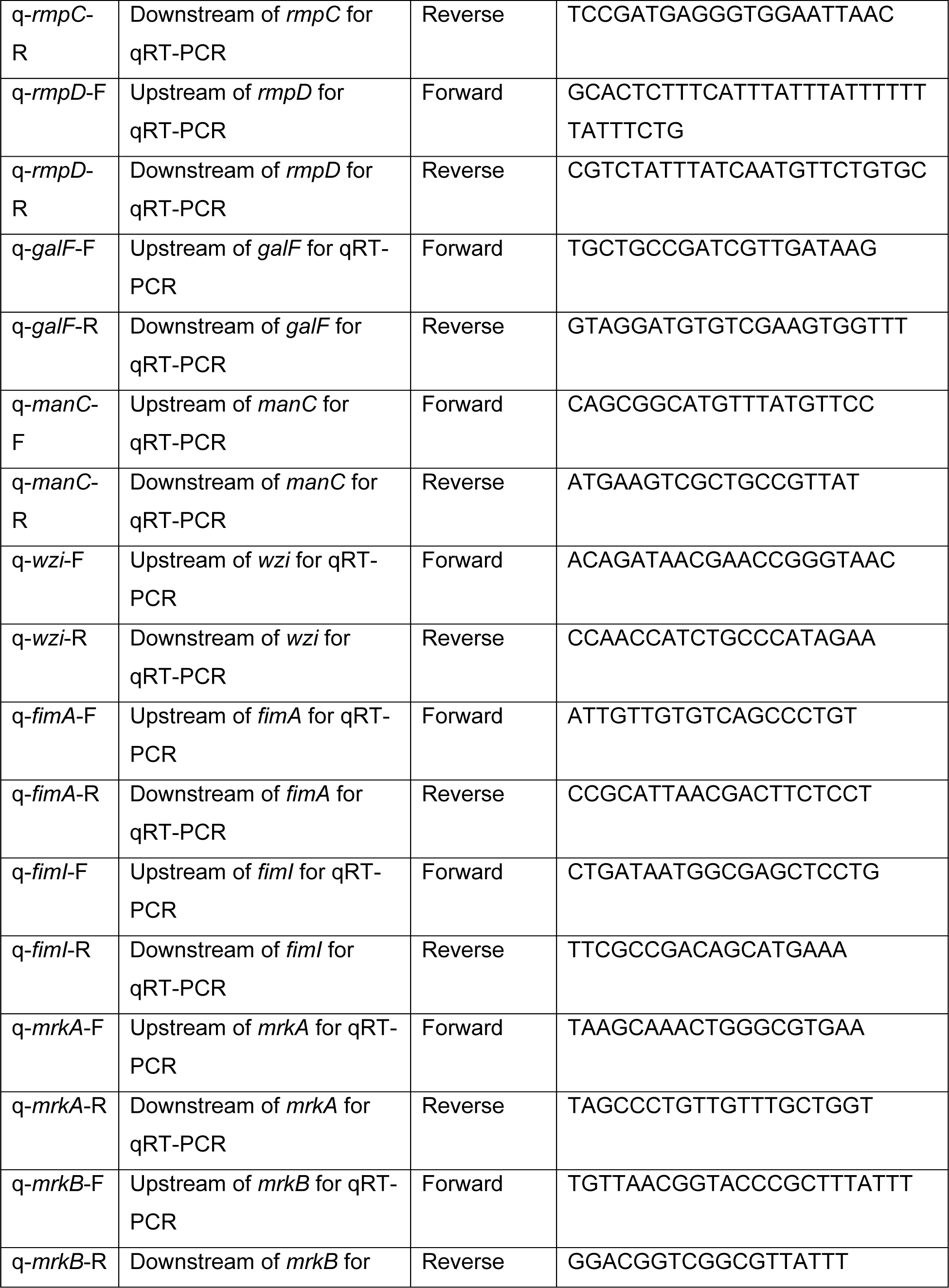
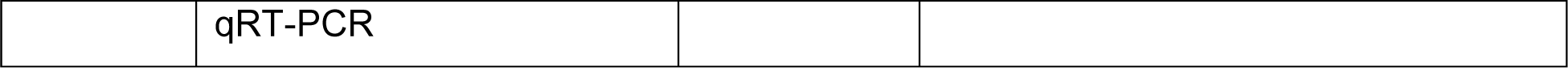
List of primers used in the study.

